# Microaerobic Copper Stress Redirects Pyruvate Metabolism and Reveals a CopL-linked Nitrogen Response in *Staphylococcus aureus*

**DOI:** 10.64898/2026.06.18.732922

**Authors:** Sean C. Brennan, Inderpreet Kaur, Daniella Spencer, Jo Purves, Hannah R. Sampson, Julian M. Ketley, Joan A. Geoghegan, Peter W. Andrew, Kevin J. Waldron, Julie A. Morrissey

## Abstract

Copper is both an essential enzyme cofactor and an antimicrobial agent deployed by the host immune system to eradicate micro-organisms. The epidemic community-acquired methicillin-resistant *Staphylococcus aureus* (CA-MRSA) lineage USA300 carries mobile genetic elements that encode *copX*/*B* and *copL*, conferring hyper-resistance to copper, but the role of CopL beyond extracellular copper sequestration remains unclear. We have combined RNA sequencing with targeted metabolite assays under microaerobic conditions, more reflective of host environments, to define key copper induced responses in WT and *copL* mutant strains.

Subinhibitory copper exposure in microaerobic conditions triggered a distinctive transcriptional response across multiple biological functions. Unlike previous studies, copper exposure did not induce an oxidative stress response. Instead, classical copper resistance, teichoic acid modification, immune-evasion factors and core metabolic genes were induced, while genes for stress responses, metal homeostasis and virulence were repressed. Gene set enrichment analysis (GSEA) identified regulation by multiple global regulators, e.g. SigB, CodY, CcpA, Agr and Sae.

Copper exposure affected metabolism, redirecting pyruvate flux toward acetoin and lactate production rather than acetate, accompanied by coordinated shifts in TCA cycle and amino acid pathways, including glutamate accumulation. Inactivation of *copL* revealed a distinct adaptive response, with strong induction of nitrogen metabolism genes and nitrite reduction. Together, these data show that copper functions as a regulatory signal, triggering coordinated transcriptional and metabolic remodelling that potentiates *S. aureus* fitness in the host.

## Introduction

Copper is an important constituent of biological systems, but there are substantial gaps in our knowledge of how this metal operates in essential processes. Elucidation of the full picture of the mode of action of copper is not only of intrinsic value, but also offers new routes to improve health. An under-researched aspect is the effect of copper on the behaviour of microbial pathogens, beyond its direct toxicity.

Copper is an essential micronutrient required by most organisms, but due to its toxicity at higher concentrations, it is also used as an antimicrobial in healthcare, agriculture and by mammalian hosts. In the human body, copper is distributed across multiple niches to maintain normal cell function and is strictly regulated to avoid both copper deficiency and self-toxicity^1^.

Copper’s importance stems from its ability to cycle between reduced (Cu^+^) and oxidised (Cu^2+^) states, making it a catalytic co-factor for numerous metalloenzymes. Excess copper can cause toxicity through DNA damage, generation of reactive oxygen species (ROS), and protein mismetallation^2–4^. The immune system can exploit copper toxicity as an antimicrobial *in vivo* through trafficking copper ions to infection sites and within macrophage phagolysosomes to eradicate infectious microorganisms^5–8^.

To counteract the antimicrobial activity of copper, bacteria have evolved dedicated resistance mechanisms to regulate toxic concentrations, primarily through efflux or sequestration^9–13^.

For example, *Staphylococcus aureus*, an opportunistic pathogen that can colonise diverse host sites, utilise the conserved copper efflux protein CopA and the copper chaperone CopZ within their core genome for copper resistance^14–17^.

Importantly, community acquired, and healthcare-associated epidemic methicillin resistant *Staphylococcus aureus* (CA-MRSA and HA-MRSA respectively) strains have acquired additional mechanisms that confer copper hyper-resistance, and subsequently, increased resistance to macrophage killing^12,13^. The copper-hyper resistance of CA-MRSA lineages (USA300-NAE and -SAE) was acquired through mobile genetic elements (MGEs), termed the arginine catabolic mobile element (ACME) and the copper and mercury resistance element (COMER)^18,19^. Both MGEs have a conserved locus, which harbours the *copXL* operon^12,20^. This operon encodes for a P_1B-3_-type ATPase copper efflux transporter (CopX) and a surface lipoprotein (CopL)^12^. CopX is proposed to efflux copper similar to the biological function of CopA. For CopL, the precise is unknown, although it can bind extracellular copper, which is proposed to limit intracellular accumulation and subsequent antibacterial activity^12,20^.

Other *S. aureus* lineages (CC22, CC30 and CC398) also harbour additional copper resistance loci from MGEs. These strains have acquired the *copBmco* operon, which encode another P_1B-3_-type copper efflux transporter, CopB, and a multicopper oxidase (Mco)^13,21^. All these copper resistance genes, both within the core genome and those that have been horizontally acquired, are regulated by the conserved copper-dependent repressor CsoR^12,20,21^.

The acquisition of these additional copper loci in *S. aureus* has resulted in reduced rates of killing of these strains by macrophages, highlighting the importance of copper homeostasis in innate immune defences^12,13^. Furthermore, the acquisition results in increased fitness in whole human blood, lung, skin and urinary tract infections, providing further compelling evidence of the significance of copper resistance in the success of the USA300 epidemic strain^8,22^.

Copper toxicity has been linked to oxidative stress via a Fenton-like reaction between oxygen, hydrogen peroxide and copper which produces hydroxyl radicals^23^. However, infection sites, including within phagosomes and biofilms, are often microaerobic or hypoxic, where oxygen availability and copper reactivity may differ significantly^24–27^. And yet, most studies of *S. aureus* copper responses have been conducted in well aerated, shaking cultures, where oxygen transfer is rapid and sustained. These culture conditions induce conserved patterns of oxidative stress pathways, protein damage and subsequent protein repair systems^22,28,29^. We hypothesised that in microaerobic conditions a completely different set of genetic responses would be seen in the presence of copper.

Here we present the first combined transcriptomic and phenotypic analysis of CA-MRSA USA300 grown under microaerobic conditions in a defined medium (RPMI-A) in the presence of subinhibitory copper concentrations. Transcriptomic analysis revealed no observed induction of oxidative stress or misfolded protein stress genes, even at copper concentrations conducted in previous studies under aerobic conditions^22,28,29^. In contrast, our data show that copper exposure induces a distinctive, adaptive transcriptional response, altering pyruvate and amino acid metabolism alongside differential expression of virulence and cell envelope genes. Additionally, comparative profiling of a *copL* mutant reveals that copper responsive regulation extends beyond detoxification and provides links with nitrogen and carbon metabolism. These findings suggest that in *S. aureus,* copper is a key regulatory signal during infection, inducing physiological adaptations that are advantageous for survival in different niches within the host.

## Results

### Global adaptive transcriptional response of *S. aureus* to copper exposure under microaerobic conditions

To define the *S. aureus* USA300 copper adaptive response in microaerobic conditions, the wild-type (JE2 WT) and a JE2 *copL::TNT* mutant strains (*copL* herein) were grown statically in RPMI-A (copper free) in 5% CO_2_. Measurement of dissolved oxygen in the static bacterial growth medium in 5% CO_2_ was compared to shaking conditions under a normal atmoshpere and provided strong confirmation that microaerobic conditions were reached (Figure S1 A and B). The markerless *copL::TNT* strain was constructed by allelic recombination using pTNT to replace the erythromycin resistance gene in the *ΦNΣ* transposon^30^. The strain was then confirmed by PCR. Additionally, RT-qPCR of *copX* and *copL* transcripts showed that, within the *copXL* operon, only *copL* transcription was disrupted (Figure S2).

To characterise the global transcriptional response, both wild-type (WT) and *copL* mutants were grown either with or without added 100 μM CuCl_2_ in microaerobic conditions. These conditions did not cause any significant growth inhibition in either the WT type or *copL* mutant strain. RNA was extracted at mid-exponential phase, and differential gene expression was analysed using the DESeq2 pipeline. Genes were considered significantly altered in expression if they demonstrated a ≥ 1.0 or ≤ −1.0 log_2_ fold change and an adjusted p-value ≤ 0.01. Principal component analysis (PCA) of all 12 samples was performed, revealing clear clustering by treatment rather than strain, indicating that copper exposure was the dominant driver of transcriptional variation (Figure 1A). One *copL* control sample clustered separately from its group, but the overall separation between copper-treated and control samples was nonetheless robust. In total, there were 181 genes which were differentially expressed in response to the addition of 100 μM CuCl_2_ (Figure 1B); 65 genes were significantly upregulated, whilst 116 were downregulated. The copper-regulated transcriptome indicated regulatory shifts across multiple biological processes (Figure 1C).

**Figure 1:**
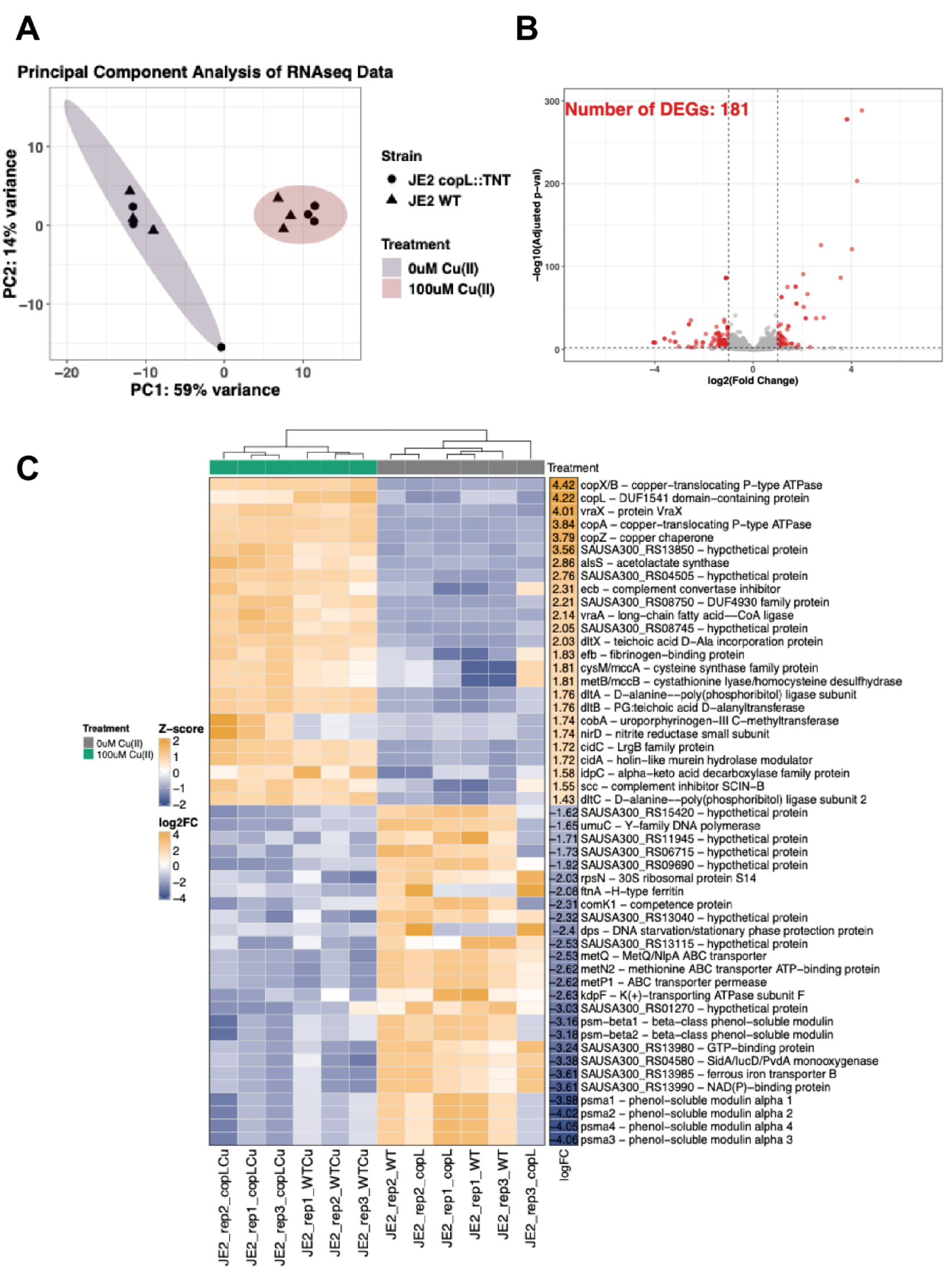
Global transcriptional response of *S. aureus* USA300 WT and *copL* mutant to subinhibitory copper exposure. (A) Principal component analysis (PCA) of the RNA-seq data set. Sample clustering was primarily driven by copper, regardless of the strain background, except for one *copL* control replicate. WT samples are depicted as triangles and *copL* samples are depicted as circles. Copper treatment is shaded appropriately to visualise clustering. (B) Volcano plot depicting differential gene expression of *S. aureus* in response to 100 μM versus 0 µM CuCl_2_ controls independent of strain. Each point represents a single gene. Genes that meet the filtering criteria, with log_2_ fold change of > 1 or < −1 and adjusted p-value < 0.01, are coloured red (differentially expressed). Horizontal and vertical dashed lines mark the filtering conditions of adjusted p-value and log_2_ fold change respectively. (C) Heatmap of the top 50 differentially expressed genes (25 most upregulated and 25 most downregulated) across all 12 RNA-seq samples. Expression values are row-wise Z-scores of regularised log-transformed counts (rlog) and columns are clustered by treatment. Rows represent the individual gene annotated by the gene name or locus tag and product name.

To explore these changes at the molecular pathway level, gene set enrichment analysis (GSEA) was performed using curated gene sets from KEGG and previous large RNAseq computational studies (Figure 2A)^31,32^. This approach evaluates the coordinated behaviour of groups of functionally related genes, rather than individual genes, to identify biological pathways or processes that were most affected by copper.

**Figure 2:**
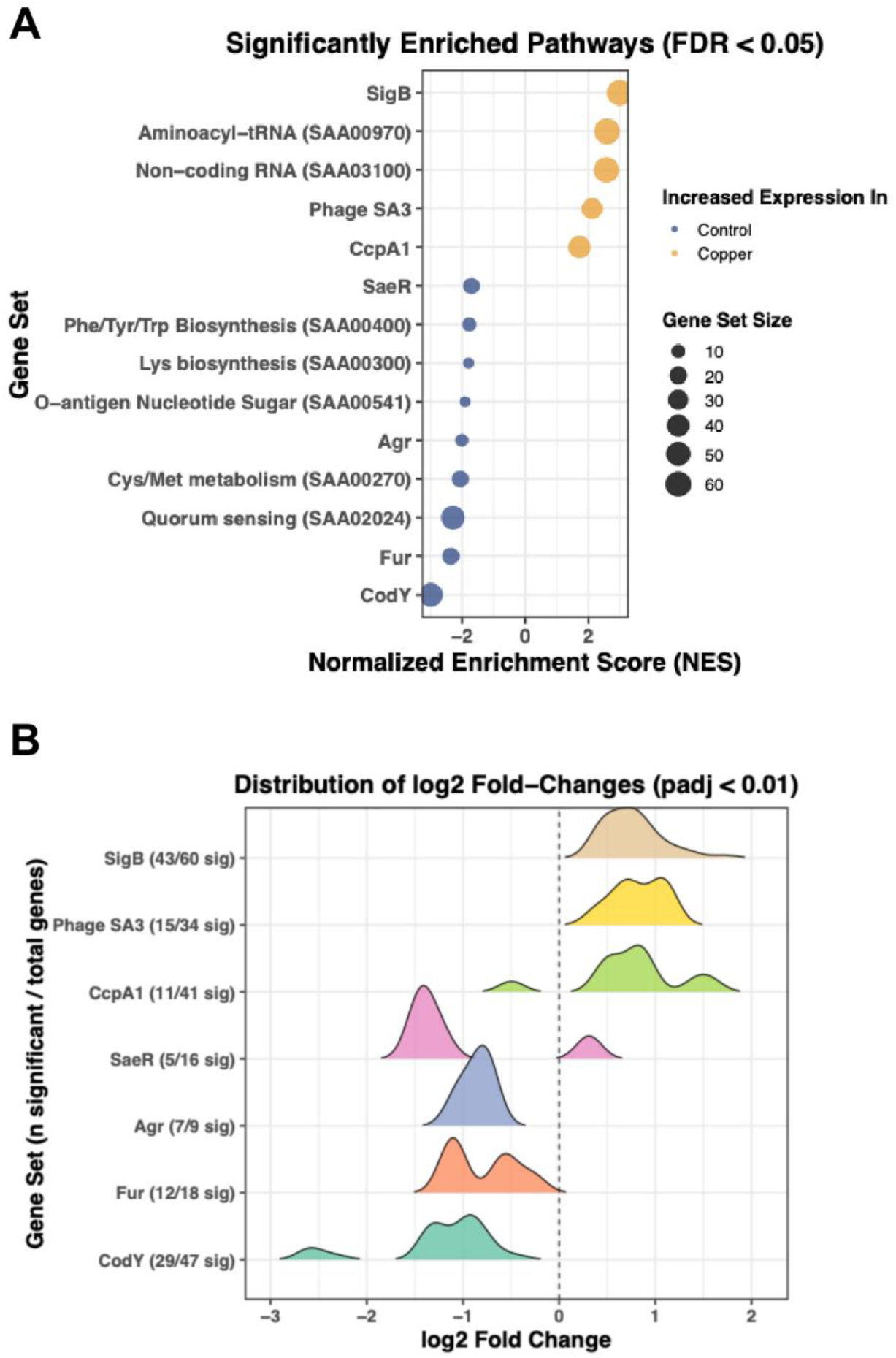
Gene set enrichment analysis (GSEA) reveals distinct changes in regulatory systems and pathways. (A) Gene set enrichment analysis (GSEA) dot plot showing significantly enriched pathways (FDR <0.05) under 100 µM versus 0µM CuCl_2_ control. Each dot represents a gene set. The x axis is the normalised enrichment score (NES) generated from the GSEA, and the size of each dot reflects the number of genes in that set that were differentially expressed. Dots coloured orange indicate sets whose genes were collectively upregulated under copper exposure, while dots coloured blue indicate sets whose genes were collectively downregulated under copper exposure. The gene sets are ordered on the y-axis by descending NES. (B) Ridge plot of selected regulator-defined gene sets, showing the distribution of log_2_ fold change values (p_adj_ <0.01) for each set. The y-axis lists the seven enriched transcriptional regulators. Labels indicate the number of genes in that set meeting the DeSeq2 filtering criteria over the total number of genes within the set. Each ridge shows the density of log_2_ fold change values across the significantly expressed genes. Peaks shifted to the right represent a collective upregulation in response to 100 µM Cu and peaks to the left represent downregulation. A vertical line at 0 indicates no change.

Of the upregulated gene sets, the regulon of the alternative sigma factor SigB was the most strongly enriched pathway (Figure 2B). This indicates that SigB plays a role in coordinating adaptation to copper exposure. Other significantly enriched pathways were related to phage SA3, aminoacyl-tRNA biosynthesis and non-coding RNAs (Figure 2A), suggesting altered protein synthesis and extensive transcriptional reprogramming in response to copper. The carbon catabolite protein (CcpA) regulon, which regulates transcription by sensing changes to primary carbon sources such as glucose, was also significantly enriched (Figure 2A-B)^33^.

Simultaneously, the downregulated gene sets were dominated by those for amino acid metabolism regulated by the metabolic regulator CodY, which senses both branched chain amino acids and GTP (Figure 2A-B)^34^. The downstream targets of the iron response regulator, Fur, were also repressed which indicates tight coordination of metal homeostasis to limit intracellular metal ion toxicity. As previously reported for *S. aureus* SH1000, Agr and Sae targets were copper repressed, including phenol-soluble modulins, nuclease and Spl proteases (Figure 2A-B)^28^. We conclude that subinhibitory copper drives a coordinated and complex transcriptional response under microaerobic conditions, which appears to be controlled by multiple large regulatory networks, influencing metabolism, metal homeostasis and virulence.

### Conserved changes in copper efflux, cell envelope remodelling and virulence

The individual genes that were differentially expressed in response to copper exposure in microaerobic conditions are presented within the supplementary data, with genes of unknown functions excluded for clarity (Supplementary tables 1 and 2). Consistent with previous analyses of aerobic cultures^28,29^, the core copper resistance response was strongly induced in microaerobic conditions. All copper efflux loci, the copper efflux genes *copA* and *copX,* the lipoprotein *copL* and the copper chaperone *copZ*, were the most strongly induced transcripts within the RNAseq analysis (≥ 3.79 log_2_-fold change). In addition to genes for detoxification, copper induced the transcription of cell envelope genes. The *dltABCD* operon and *dltX*, which is responsible for the D-alanylation of teichoic acids, and *mprF*, which incorporates lysyl-phosphatidylglycerol into the membrane, were all significantly upregulated (1.03-2.03 log_2_-fold). Induction of copper and cell envelope stress responses has also been observed under several growth and media conditions, suggesting these are conserved features of the copper stress response^28,29^.

Copper repression of virulence-associated genes, nucleases, toxins and immune evasion genes (*nuc, splB, psmβ1*, *psmβ2*, *psmα1-3, chs*, *scpA*, *scn*, and *flr*) (−1 - −4 log_2_-fold) was also conserved between WT and *copL* strains and between microaerobic and aerobic growth conditions^28,29^. In contrast to aerobic conditions, genes involved in immune evasion were upregulated in response to copper, e.g. *ecb* and *scc*, which encode complement inhibitors, *efb,* a fibrinogen binding protein, *hlgA*, encoding a haemolysin, and *vraX,* associated with resistance to complement^35^.

Together, these data show that copper induction of copper efflux and cell envelope stress responses as well as repression of virulence toxin and proteases genes is conserved between different *S. aureus* lineages, growth and media conditions, demonstrating that there is a conserved core copper stress response^28,29^.

### Differential changes in oxidative, iron and iron-sulphur Fe-S stress responses

Subinhibitory copper exposure under microaerobic conditions did not lead to induction of oxidative stress responses, in contrast to previous studies performed under aerobic conditions^28,29^. In fact, under microaerobic conditions several key oxidative stress, iron uptake, competence and DNA repair genes were significantly downregulated by exposure to copper. This included the alkyl hydroperoxide reductase system (*ahpF* and *ahpC*), iron uptake transporters (*isdBCDEFG)*, siderophore biosynthetic genes (*sbnCDEFGH*), and stress regulators *comK* and *lexA*, *dps*, and a YolD family protein, regulated by LexA^36,37^. Furthermore, there were no significant changes in the genes encoding for iron-sulphur (Fe-S) repair systems. These genes are typically induced in aerobic conditions because Fe-S proteins are sensitive to copper-induced oxidative damage^3,38^. RT-qPCR of oxidative stress gene expression confirmed that there was no induction of these genes by copper under microaerobic conditions (Figure S2).

### Copper induces transcriptional shifts in genes for pyruvate and amino acid metabolism

Metabolism was differentially regulated in response to copper under microaerobic conditions. In aerobic conditions, due to copper induced damage to the glycolytic enzyme glyceraldehyde-3-phosphate dehydrogenase A (GapA/GAPDH), glycolytic genes are induced and there is redirection of carbon flux towards the pentose phosphate pathway (PPP)^22,29^. In microaerobic conditions these genes were not transcriptionally altered by copper. In contrast, the expression of pyruvate metabolism genes was induced. Alternatively, the *cidABC* operon (log_2_-fold 1.40-1.73), associated with pyruvate oxidation to acetate and *alsS* (log_2_-fold 2.86), which diverts pyruvate to acetoin via *alsD*, were significantly upregulated^39^.

Genes for amino acid metabolism was also altered in response to subinhibitory copper exposure under microaerobic conditions. The anabolic arginine and cysteine biosynthesis genes were upregulated, but conversely, there was significant downregulation of branched-chain amino acids, methionine and histidine biosynthetic genes, many of which are regulated by CodY^40^. Together, these data support the idea that copper acts as a signal, triggering transcriptional and metabolic changes that are differential dependent on oxygen levels.

### Copper exposure shifts pyruvate metabolism under microaerobic conditions

To investigate the effect of copper on pyruvate metabolism, *S. aureus* WT and the pyruvate oxidase (*cidC*) and acetolactate synthase (*alsS*) transposon insertion mutants (*cidC::Tn* and *alsS::Tn*) were utilised. Each strain was grown microaerobically to mid-exponential phase, with or without added 100 μM CuCl_2_ and the abundance of the secreted end-products of the pyruvate pathways: pyruvate, acetoin, acetate and lactic acid was measured (Figure 3)^41^. D-glucose utilisation was also calculated by quantifying glucose in culture supernatants as previously described^41^.

**Figure 3:**
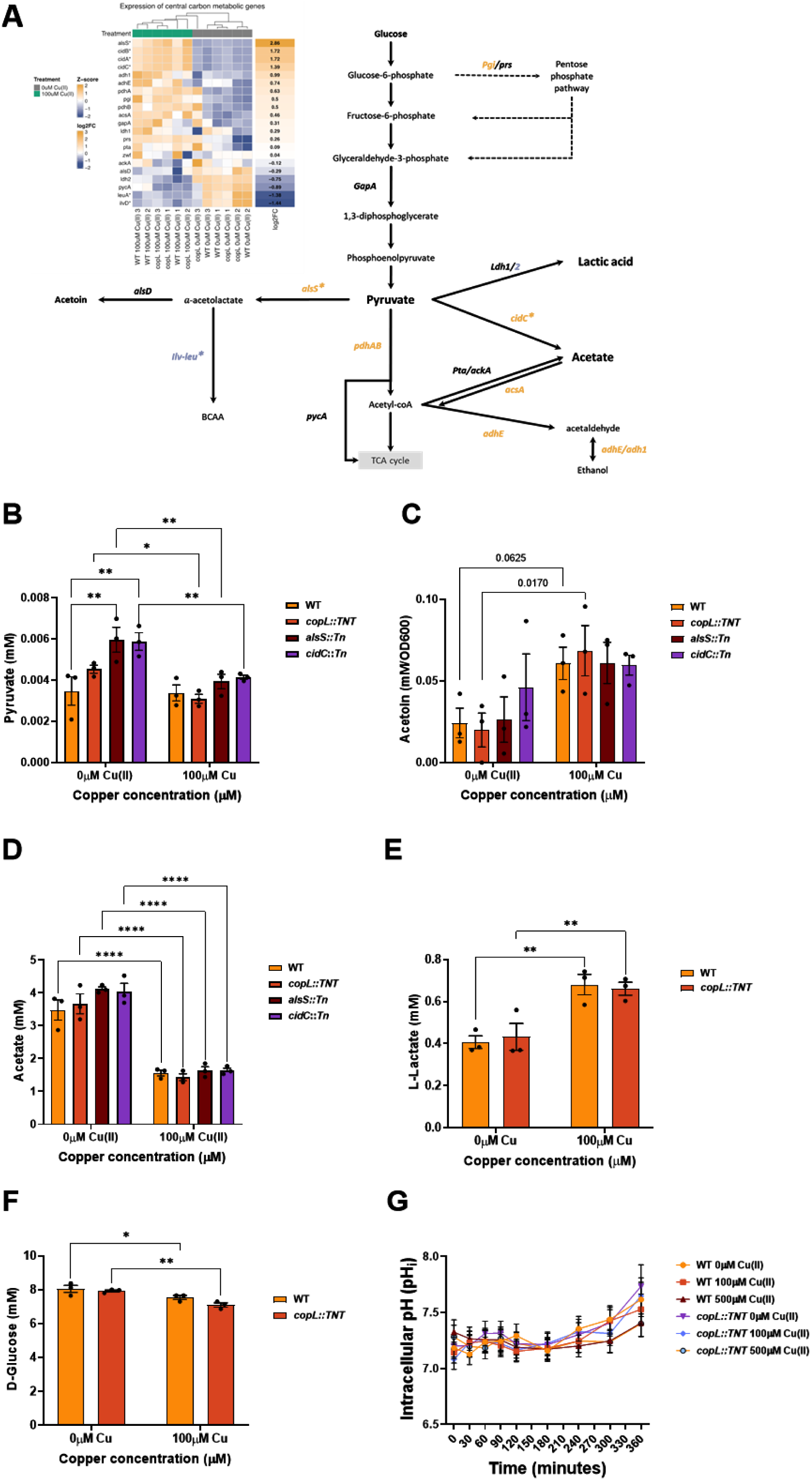
Copper exposure alters pyruvate metabolism and glucose consumption in *S. aureus*. (A) Schematic overview of glycolysis and downstream pyruvate metabolic pathways in *S. aureus.* Gene expression data are overlaid as a heatmap showing row-wise Z-scores across all RNA-seq samples. Genes that are significantly differentially expressed (adjusted p ≤ 0.01; Log_2_ fold change ≥ 1 or ≤ −1) in response to 100 µM CuCl_2_ are marked with an asterisk (*). Genes upregulated are shaded orange whilst those downregulated are shown in blue. Arrows next to certain metabolites depict significant extracellular concentrations in the WT and *copL* mutant, depicted in B-F. (B-F) Concentrations of key pyruvate metabolism end-products were quantified in culture supernatants of *S. aureus* USA300 JE2 WT, *copL*, *alsS*::Tn, and *cidC*::Tn mutants, grown statically at 37 °C and 5% CO_2_ in RPMI-A, with or without 100 ***u***M CuCl_2._ Metabolite concentrations for (B) pyruvate, (C) acetoin and (D) acetate were normalised to OD600. Concentrations for (E) lactic acid and (F) D-glucose were also quantified. Error bars for each metabolite represent ± the standard error of the mean (SEM) from at least 3 biological replicates. Significance between conditions was determined using a two-way ANOVA test with a Tukeys multiple comparisons test (**** = P ≤ 0.0001, ** = P ≤ 0.01, * = P ≤ 0.05). (G) Intracellular pH was measured in WT (circles) and *copL* (squares) strains were grown up to 6 hours under microaerobic conditions supplemented with 0, 100, or 500 µM CuCl₂ and then stained with CFDA-SE. Fluorescence ratio (490/444 nm) was used to interpolate intracellular pH from a calibration curve (Supplementary Data). Data represent the mean ± SEM from at least 3 biological replicates.

In the absence of copper, the supernatants of the *alsS* and *cidC* mutants exhibited significantly higher concentrations of extracellular pyruvate compared to the WT (P < 0.01, Figure 3B), suggesting blockage of pyruvate processing resulting in the metabolite being excreted. In the presence of subinhibitory copper, the concentration of pyruvate in WT supernatants was statistically unchanged (P>0.05). However, pyruvate concentrations in supernatants of *copL, alsS* and *cidC* cultures were decreased significantly compared to controls (P < 0.01). These data suggest that the observed increased expression of both *alsS* and *cidC* plays a role in pyruvate metabolism in response to copper exposure.

Acetoin levels in the supernatant of all strains in microaerobic conditions were similar (P > 0.05; Figure 3C). All four strains appeared to exhibit increased acetoin in 100 μM copper, but only the *copL* mutant culture was determined to be statistically significant (P < 0.05). Conversely, extracellular acetate was significantly decreased in all strains when grown in the presence of subinhibitory copper (P < 0.0001), despite significant upregulation of *cidC* (Figure 3A and 3D). Overall, acetoin production changes were modest relative to the stronger and consistent acetate phenotype. The lower acetate concentrations suggest either preferential synthesis of acetoin over acetate or acetate consumption in microaerobic, copper-exposed cells into acetyl-CoA.

An additional by-product of pyruvate metabolism, lactic acid, was also quantified in cultures of the WT and *copL* mutant (Figure 3E). Interestingly, despite no transcriptional evidence of induction of the *ldh* gene, both *S. aureus* strains produced significantly increased concentrations of lactic acid in the supernatant compared to growth in the absence of copper (P < 0.01).

To determine if copper altered glucose utilisation in microaerobic conditions, D-glucose consumption was also quantified in the WT and *copL* mutant strains (Figure 3F). There were significant decreases in D-glucose consumption by both WT and *copL* strains when grown in subinhibitory copper concentrations (P < 0.05; P < 0.01, respectively) compared to culture in the absence of copper. Together, these data show that exposure to copper under microaerobic conditions increases the utilisation of glucose and causes a rebalance of pyruvate metabolism resulting in a consistent decrease in extracellular acetate, accompanied by increased lactate production and an acetoin increase.

It was previously described that the catalysis of pyruvate to acetoin via the *alsS* operon utilises intracellular protons and can protect the cell from intracellular pH acidification during growth on glucose^42^. To test this concept, the intracellular pH in the WT and *copL* mutant grown microaerobically in the presence of 0, 100 or 500 µM copper and intracellular pH was measured using the 5-(6)-carboxyfluorescein diacetate N-succinimidyl ester (cFDA- SE) fluorescent probe. The intracellular pH of the WT or *copL* mutant was not altered over time in any concentration of copper (P > 0.05; Figure 3G), showing that intracellular pH is controlled during glucose consumption and copper stress.

### Copper induces SigB- and CidR-dependent control of pyruvate metabolism genes

Following the demonstration that copper induces the *als* and *cid* operons to redirect pyruvate flux, the regulatory factors behind the metabolic rerouting were investigated. The alternative sigma factor SigB, along with the CidR transcriptional activator, are involved in the regulation of the *als* and *cid* operons in conditions of excess glucose ^39,43,44^. Furthermore, the SigB regulon is the most differentially regulated in response to copper in this study. Thus, we hypothesised that SigB and CidR play a role in *S. aureus* USA300 copper-responsive gene regulation.

To test this hypothesis, *sigB::Tn* and *cidR::Tn* mutants were constructed into a clean USA300 JE2 background, and growth assays confirmed that neither mutation altered *S. aureus* growth at 100 μM or 500 μM (inhibitory) copper concentrations compared to the wild type (Figure S3). Next, WT, *sigB* and *cidR* strains were grown microaerobically with or without added 100 μM CuCl_2_ and comparative gene expression of the copper efflux genes (*copA* and *copL*) and pyruvate metabolism genes (*alsS* and *cidB*) were quantified by RT-qPCR. Analysis of the expression of the copper homeostasis genes *copA* and *copL* revealed no difference between the *cidR* or *sigB* mutants and the WT, in both the absence or presence of copper. These data demonstrate that neither regulator influences copper homeostasis genes under these conditions (P > 0.05; Figure 4A). Contrastingly, in 100 μM copper, both *alsS* and *cidB* expression was significantly upregulated in the WT strain, consistent with the RNAseq data (P ≤ 0.0001; P ≤ 0.05 respectively; Figure 4B). In both the *sigB* and *cidR* mutants, *alsS* was not induced by copper (P >0.05 vs. WT control; Figure 4B) and was significantly lower than in WT cells after copper treatment (P < 0.0001), indicating that both regulators are required for *alsS* gene activation in copper environments. Similarly, *cidB* induction in response to copper was abolished in both mutants, with the *sigB* strain showing significant decreases under both experimental conditions in comparison to the WT (P < 0.0001; Figure 4B).

**Figure 4:**
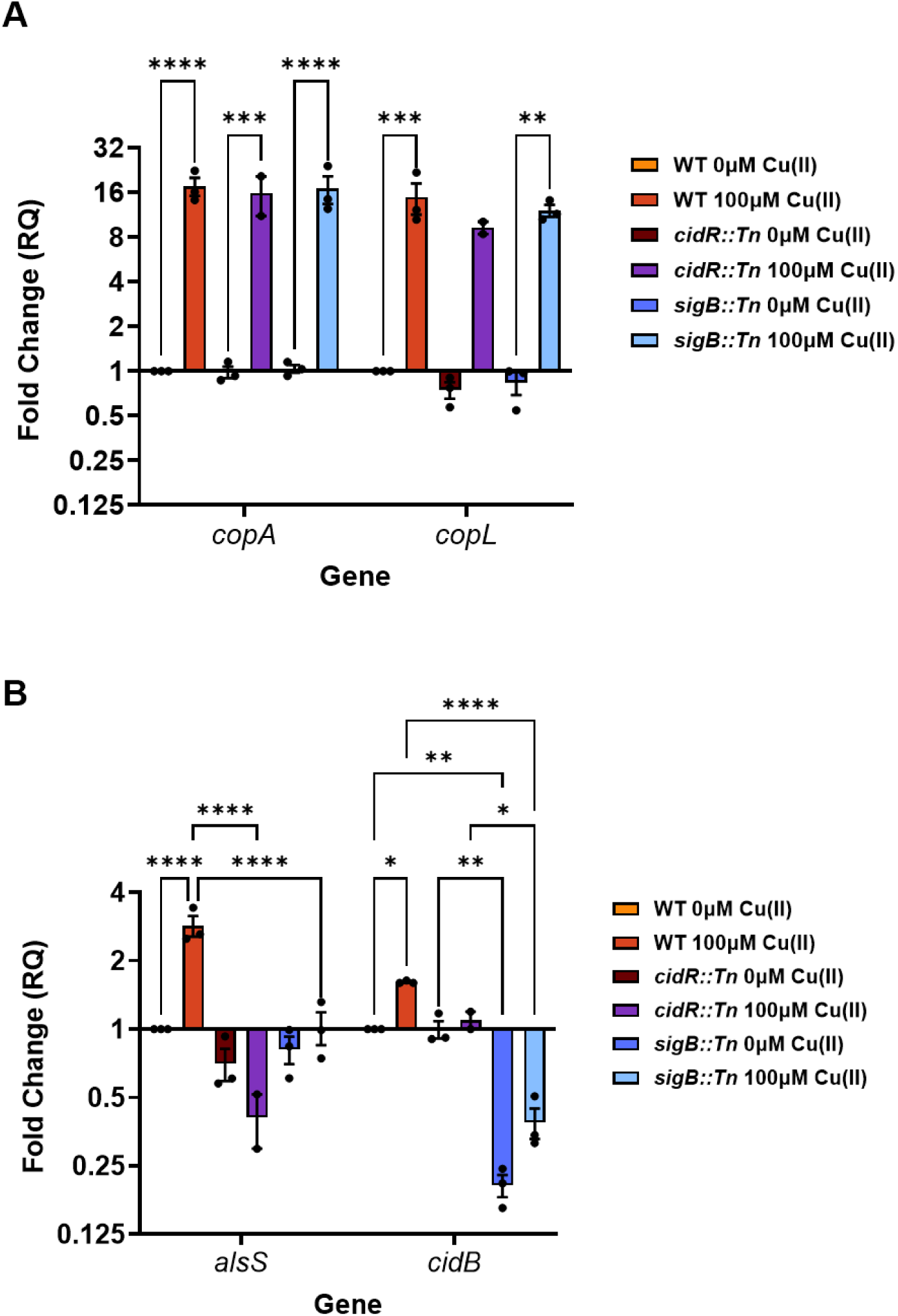
Transcriptional regulation of *copA, copL, alsS* and *cidB* in response to copper in regulatory mutants. (A-C) Relative expression of *copA, copL* (A), *alsS*, and *cidB* in the *SigB*::Tn (B) and *cidR*::Tn (C) mutant strain was assessed by RT-qPCR following exposure to 0 or 100 µM CuCl_2_ during mid-exponential growth in RPMI-A at 37 °C + 5 % CO2. Expression values were calculated using the ΔΔCt method, normalised to the endogenous control *gyrB*, and expressed relative to the wild-type USA300 JE2 strain under no copper exposure (RQ = 1). Mid-exponential phase at 37 °C + 5 % CO2 in RPMI-A supplemented with no copper or 100 µM CuCl2 was measured. Results were normalised to OD600nm. Error bars represent the ± standard error of mean (SEM) from at least 3 biological replicates. Significance between conditions was determined using a two-way ANOVA test with a Tukeys multiple comparisons test (*P ≤ 0.05; **P ≤ 0.01; ***P ≤ 0.001; ****P ≤ 0.0001; ns = not significant)

### Copper exposure increases TCA activity and intracellular glutamate concentration

Building on the observed rebalancing of pyruvate metabolism, changes in other central metabolic pathways were explored, namely the TCA cycle and its integration with amino acid metabolism. PCA and clustering analyses (Figure 1A) revealed a consistent increase across copper-exposed samples of the *gltA, acnA, icd, sucCD, sdhABC* and *fumC* genes for the TCA cycle, as well as *glnA, rocA/pruA, argGH*, and *rocF* for glutamate/arginine metabolism (Figure 5A).

**Figure 5:**
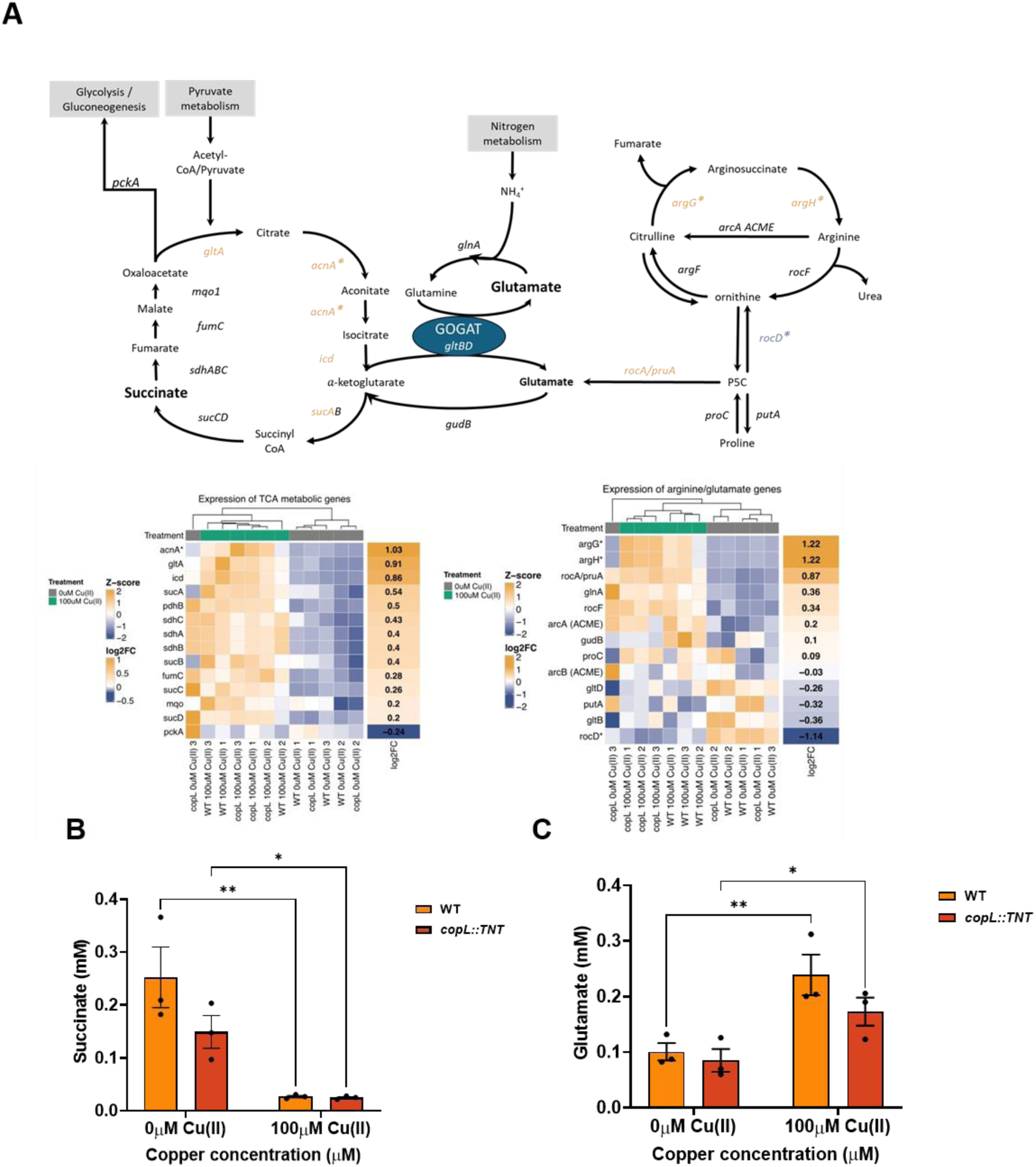
Copper alters glutamate synthesis and TCA cycle activity. (A) Schematic overview of the tricarboxylic acid (TCA) cycle and the argininosuccinate biosynthesis pathways in *S. aureus*. Gene expression data are overlaid as heatmaps, showing row-wise Z-scores across all RNA-seq samples. Genes significantly differentially expressed in response to 100 µM CuCl₂ (adjusted p ≤ 0.01; log₂ fold change ≥ 1 or ≤ –1) are marked with an asterisk (*). Genes showing upregulation under copper exposure are shaded orange, and those downregulated are shaded blue. Heatmaps are grouped by pathway, highlighting transcriptional shifts in central metabolism and amino acid biosynthesis under copper stress. (B) The concentration of succinate in the lysates of mid-exponentially growing USA300 JE2 WT and *copL* mutant grown to mid-exponential phase at 37 °C + 5 % CO2 in RPMI-A supplemented with no copper or 100 µM CuCl2 was measured. Results were normalised to OD600nm. Error bars represent the ± standard error of mean (SEM) from at least 3 biological replicates. Significance between conditions was determined using a two-way ANOVA test with a Tukeys multiple comparisons test (NS = no significant difference, **** = P ≤ 0.0001). (C) The concentrations of intracellular glutamate was measured from mid-exponentially growing *S. aureus* USA300 JE2 and *copL* mutant at 37°C + 5% CO2 in RPMI-A -/+ 100μM CuCl2 using the Glutamine/Glutamate-Glo Assay (Promega). Error bars represent ± standard error of mean (SEM) from 3 independent biological replicates. Significance was determined using two-way ANOVA Tukey’s multiple comparisons test.

To determine intracellular metabolite concentrations, WT and *copL* mutant cells were grown microaerobically with or without 100 μM added CuCl_2._ The measurement of succinate and glutamate were conducted using colorimetric and luminescence assays respectively.

Succinate concentrations were measured using cell lysates from rapid homogenisation. To measure intracellular glutamate, metabolism was stopped using a HCl inactivation solution. Compared to control conditions, a significant decrease in the intracellular concentration of succinate in both the WT and *copL* mutant in response to copper exposure was observed (P < 0.0001; Figure 5B), with no significant difference between the wild type and mutant strains (P > 0.05).

The concentrations of intracellular glutamate in the absence of copper were the same in the cultures of the WT and *copL* mutant strain at ∼20 μM. In response to copper, glutamate concentrations increased significantly in both the WT and *copL* strains (P = <0.01 and <0.05, respectively; Figure 5C). To investigate whether elevated glutamate contributes to copper tolerance, *S. aureus* WT and *copL* were grown under inhibitory copper concentrations (1 mM) with increasing concentrations of L- or D-glutamate or proline, which is a substrate for glutamate production. D-stereoisomers of the amino acids were also included as they are not primarily metabolised by bacteria. Bacterial growth was rescued with supplementation with both amino acids compared to copper exposure alone, regardless of the stereochemistry (Figure S5). The data show that glutamate and proline metabolism contribute to copper tolerance potentially via extracellular chelation.

### CopL promotes enhanced nitrate/nitrite reduction in response to copper

Throughout this study, both the WT and *copL* mutant samples were analysed to define the core transcriptional response to 100 μM CuCl_2_ under microaerobic conditions. To determine whether CopL has unknown functions in the copper response, a Spearman correlation was performed comparing RNAseq transcriptional log_2_ fold changes in the WT ± copper versus *copL* ± copper (Figure 7A). The correlation between the treatments was strong demonstrating that the transcriptional profiles were very similar (R = 0.81, P < 2.23-16) (Figure 6A).

**Figure 6:**
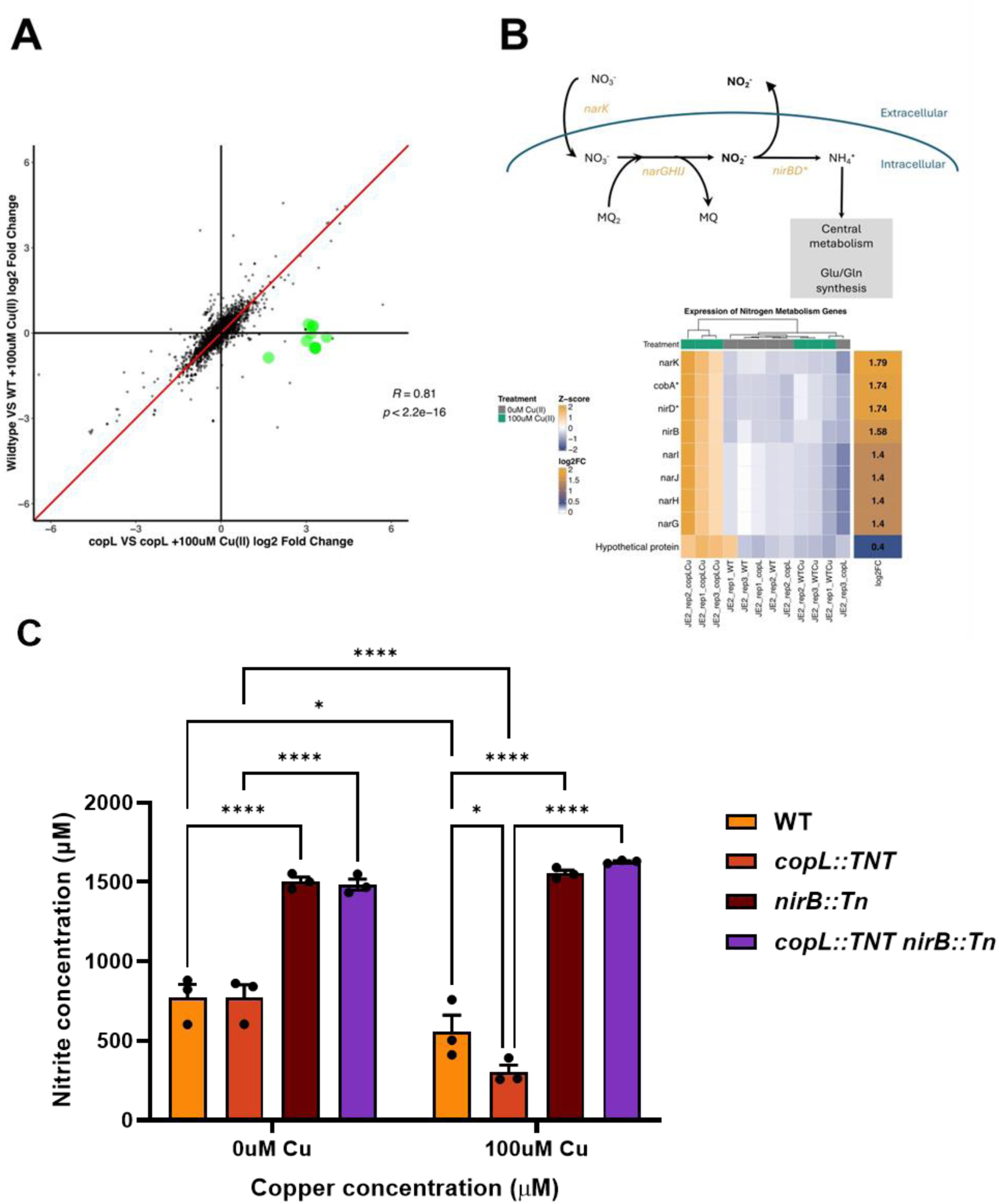
Copper induces increased rates of nitrite reduction. (A) Spearman correlation graph comparing the log2 fold changes of the WT vs. WT + 100μM Cu(II)Cl2 on the x-axis and the log2 fold changes of the *copL* mutant versus *copL* + 100μM Cu(II)Cl2 on the y-axis. The correlation coefficient (R) and corresponding p-value are displayed. The black solid lines represent log2 fold changes of 0, while the red line aids in visualizing similarities between values. Points highlighted in green indicate genes that are significantly differentially regulated in the *copL* mutant but not in the WT. Differential expression data were generated from DESeq2 analysis. (B) Schematic overview of nitrogen metabolism in *S. aureus*, including nitrate and nitrite reduction pathways. Gene expression data are overlaid as a heatmap showing row-wise Z-scores across all RNA-seq samples. Genes significantly differentially expressed in response to 100 µM Cu(II)Cl₂ (adjusted p ≤ 0.01; log₂ fold change ≥ 1 or ≤ –1) are marked with an asterisk (*). Upregulated genes are shaded orange, highlighting transcriptional adaptation of nitrogen assimilation pathways under copper stress. (C) Concentrations of nitrite was quantified in culture supernatants of S. aureus USA300 JE2 WT, copL::TNT, nirB::Tn mutants and copL::TNT/nirB::Tn double mutant grown statically at 37 °C and 5% CO2 in RPMI-A with or without 100 μM Cu(II)Cl_2_. Error bars represent the ± standard error of the mean (SEM) from at least 3 biological replicates. Significance between conditions was determined using a_2_t_5_wo-way ANOVA test with a Turkeys multiple comparisons test (**** = P≤ 0.0001, * = P ≤ 0.05).

Genes were classified as CopL-specific if they exhibited significant expression changes in the CopL comparison but not in the WT comparison. Using these criteria, we found that nitrate/nitrite reduction loci were identified as induced in the *copL* mutant in response to copper (Figure 6A-B). This included the induction of the *narGHJI* operon (encoding nitrate reductase), *nirBD/cobA* (involved in nitrite reduction to ammonia), and associated regulatory and transport genes (*nark, nirR*) and a hypothetical protein (Figure 6B). These genes fall under the control of the NreBC nitrogen regulation system, which is activated under oxygen limitation^45^. The observed upregulation of these genes specifically in the *copL* mutant suggests that, although the *copL* mutant shows the same copper response as the wild type, mutation of *copL* also triggers enhanced nitrate/nitrite reduction when grown under copper in microaerobic conditions.

To assess nitrogen metabolism activity, nitrate/nitrite reduction of the WT, *copL* and *nirB::Tn* single and double mutants was assessed. Each strain was grown to mid-exponential phase with or without 100μM CuCl_2_ in microaerobic conditions and. nitrite concentrations in supernatants were measured using a colorimetric assay.

When grown in subinhibitory copper concentrations, both the WT and *copL* produced significantly lower nitrite concentrations (P < 0.05 and <0.0001, respectively) in comparison to growth in the absence of copper, indicating faster rates of nitrite consumption by both strains (Figure 7C). Consumption of nitrite by the *copL* mutant was increased significantly compared to the WT in the presence of copper (P < 0.05), correlating with the observed increase in nitrogen metabolism gene expression in the *copL* mutant. The levels of nitrite in the supernatant of the *nirB* single and *nirB/copL* double mutant remained unchanged by exposure to copper (P>0.05) but were significantly higher in comparison to the WT and *copL* mutant under both conditions (P < 0.0001). These data demonstrate that nitrate/nitrite reduction is increased in *S. aureus* when grown in the presence of subinhibitory copper in microaerobic conditions. Together, these data suggest *S. aureus* coordinates a carbon and nitrogen metabolic response when grown statically under microaerobic conditions and subinhibitory copper, allowing energy fluxes to sustain growth in these conditions.

## Discussion

Copper is both an essential enzyme cofactor and, due to its toxicity at higher concentrations, a potent in-host antimicrobial agent, thus copper stress responses are critical for bacterial survival in a host. *S. aureus* copper stress responses are well characterised under aerobic conditions^22,28,29^, however the responses to copper in microaerobic conditions have not previously been studied. This is a critical knowledge gap because host niches and tissues have varying oxygen levels^24–27^. *S. aureus* colonises diverse host environments, including the nasopharynx, skin, blood, heart and bone, where oxygen levels vary widely and can reach microaerobic concentrations^46^. Host pathophysiology during infection also leads to significant oxygen depletion creating microaerobic environments^46^. Considering copper concentrations are known to increase at infection sites^8^, it is crucial to investigate the adaptive responses of copper under physiologically relevant microaerobic oxygen concentrations.

In this study, substantial differences are observed between microaerobic and aerobic adaptative responses to copper. A core copper-specific transcriptional response, consisting of induction of classical copper detoxification and cell envelope biosynthesis genes and the repression of toxin and protease virulence genes, is highly conserved between strains and conditions showing the core response is essential for *S. aureus* adaptation to copper in different niches. We have also uncovered an adaptive copper response that is dependent on oxygen levels. In microaerobic conditions, there is a distinct response, with expression of metabolic pathways and the oxidative stress response differing significantly to the response observed in aerobic conditions. Additionally, we found previously unknown links between copper and nitrogen metabolism. These data provide novel insight into *S. aureus* metabolic adaptation to both copper and oxygen levels critical for bacterial survival in different niches during infection.

Our data show that *S. aureus* also has a core copper-specific response that is independent of oxygen, strain and media composition. In both aerobic^22,28,29^ and microaerobic environments, the core and acquired copper genes, *copAZ* and *copXL*, are strongly induced demonstrating a highly conserved physiological adaptation. This response is also independent of whether the cell is cultured under nutrient-rich or -limited conditions. This reflects the importance of bacterial copper tolerance, consistent with copper detoxification being critical for *S. aureus* survival within macrophages and during infection^8,12,13,22^.

Additionally, there is conserved increased expression of genes for cell wall modification, *dltABCD* and *mprF*, in microaerobic and aerobic conditions^29^, suggesting that D-alanylation of teichoic acid is a central feature of the copper stress response. Both the *S. aureus dlt* operon and *mprF* have been implicated in reducing cell surface anionic charge^47,48^. Copper-mediated induction of these loci could increase cellular positive charge inducing resistance to cationic antimicrobials, potentiating staphylococcal persistence within the host. Copper-mediated upregulation of *dltABCD* operon expression also has been observed in *Enterococcus faecalis*, indicating a potentially conserved adaptation to copper exposure across species^49^.

Interestingly, we showed that there is also conserved repression of virulence-associated genes encoding for nucleases, and toxins^28,29^, that is consistent with previous studies showing copper repression of virulence genes^28^. Together, these findings suggest that elevated copper acts as a signal, that reduces virulent activity, supporting the hypothesis that copper induces the bacteria to shift to a less aggressive phenotype perhaps to prevent a strong immune response and thus potentiate colonisation in contrast to an invasive disease-causing phenotype.

This hypothesis is supported by our RNAseq data demonstrating that not all virulence factors show copper-dependent repression in microaerobic conditions. Indeed, there is induction of genes important for immune evasion, complement inhibitor genes *scc* and *ecb*, fibrinogen binding protein gene *efb* and gamma haemolysin subunit *hlgA* gene. These genes block innate complement system pathways and phagocyte killing, consistent with subinhibitory copper in microaerobic conditions inducing a *S. aureus* immune-evasive persistence phenotype^50,51^.

Although there is a core copper response irrespective of oxygen levels, *S. aureus* also has a distinct differential response in microaerobic conditions compared to aerobic conditions, with copper inducing oxidative stress pathways and protein repair systems only in aerobic conditions^28,29^. In contrast, in microaerobic conditions there is repression of oxidative stress, DNA repair, and metal homeostasis genes. Copper toxicity has been linked to Fenton-like reactions that generate reactive oxygen species (ROS) that can damage DNA, iron-sulphur clusters and proteins^3,52^, resulting in the need for induction of pathways needed to counteract the damage. The lack of copper-mediated induction of these pathways under microaerobic conditions suggest ROS formation is decreased due to reduced oxygen availability. Repression of iron uptake, a catalyst for the Fenton reaction, may also protect against copper-mediated ROS oxidative stress during microaerobic growth.

There is also a differential copper-mediated metabolic response. In aerobic conditions, copper induces changes in carbon flux to account for inactivation of the GAPDH and phosphoribosylpyrophosphate synthetase (Prs) enzymes critical for glycolysis and the pentose phosphate pathway respectively^22,29^. In contrast, in microaerobic conditions, copper alters pyruvate metabolism to induce expression of the *alsS* and *cidC* operons that typically drive pyruvate towards acetoin and acetate production respectively (Figure 3A). The change in pyruvate metabolism and the resultant alteration in extracellular metabolites has not been previously reported as part of the copper response in S*. aureus*.

End-product metabolite measurements confirm copper-mediated increases in acetoin concentrations. However, extracellular acetate concentrations decrease, suggesting either rerouting of pyruvate towards acetoin or utilisation of acetate to produce acetyl-CoA. The observed increased transcription of enzymes required for acetyl-CoA production, acetyl-coenzyme A synthetase (*acsA*) and pyruvate dehydrogenase (*pdhAB*) (Figure 3A), suggest that in microaerobic conditions, subinhibitory copper signals for increased production of both acetoin and acetyl co-A.

We propose that sub-inhibitory copper is acting as signal to induce pathways that are likely to be beneficial to *S. aureus*. The AlsS pathway has been linked with acid resistance through the consumption of protons and shuttling pyruvate from acetate and lactate production towards acetoin, a neutral metabolic end product^39,53^. Our intracellular pH measurements demonstrated that copper does not acidify the cytosol, indicating that acidification is not a copper toxicity mechanism or has been prevented. Activation of the AlsS pathway may also prime *S. aureus* to survive within macrophage phagolysosomes that are characterised by acidic pH and elevated copper concentrations^5,6,54^.

There is also copper-mediated differential expression of the TCA cycle in response to oxygen levels that could explain the need for increased acetyl co-A production. We observed increased expression of TCA cycle genes (*gltA, acnA, icd, sucCD, sdhABC* and *fumC*) and decreased intracellular succinate concentrations, suggesting increased TCA cycle activity under microaerobic conditions, that was not observed in aerobic studies. Increased TCA activity also correlates with transcriptional changes in genes for amino acid metabolism with *argGH*, for generation of fumarate from arginosuccinate, and *rocA/pruA*, for glutamate from pyrroline-5-carboxylate (P5C) and proline, both of which were significantly upregulated (Figure 5A). Together these transcriptional changes result in increased intracellular glutamate which would be beneficial for *S. aureus* because our data show glutamate chelates copper.

Many of the genes altered in the TCA cycle/glutamate/arginosuccinate network are influenced by the global carbon catabolite regulator CcpA. CcpA senses glucose concentrations and normally represses alternative metabolic pathways when glucose is abundant^32,55,56^. However, our GSEA analysis identified significant induction of the CcpA regulon, despite high glucose concentrations. Previous results revealed that copper binds to cysteine residues within CcpA, causing dissociation of CcpA from its target DNA and subsequently derepression of the regulon^57^. Although direct copper-CcpA interactions were not investigated here, the transcriptional patterns observed are consistent with a model in which copper modulates CcpA regulatory activity.

Our data also identified a novel link between copper and nitrogen metabolism under microaerobic conditions. Interestingly, copper induces expression of nitrogen metabolism genes in the *copL* mutant, but not wild type *S. aureus*. However, nitrate metabolism is increased in both the wild type and *copL* mutant, suggesting that copper induction of nitrate metabolism in microaerobic conditions is a general response and not specific to CopL. Nitrogen metabolism in *S. aureus* is regulated by the NreBC two-component system, which senses oxygen availability and contains Fe-S clusters^58^. The *copL* mutant accumulates higher intracellular concentrations of copper compared to the wild type in microaerobic conditions^12^. Therefore, elevated intracellular copper under low-oxygen environments may increase binding of copper to NreBC and induce expression of nitrogen metabolism genes in the *copL* mutant compared to the wild type. Importantly, these data create a link between copper exposure and nitrogen metabolism which has not been previously reported in *S. aureus*.

Overall, two models emerge from copper exposure in *S. aureus*, an aerobic model and a microaerobic model. Both models include a core copper response of copper detoxification, cell envelope modifications via the *dlt* operon and repression of virulence. Under aerobic conditions, we hypothesise higher levels of ROS resulting in more protein and enzyme damage. Due to this damage, *S. aureus* prioritises oxidative stress pathways to survive copper induced toxicity. Under microaerobic conditions there is limited ROS induction and therefore less protein damage and oxidative stress. Instead, *S. aureus* adapts to the copper exposure by altering central carbon metabolism via pyruvate overflow and TCA cycle activation. Through this metabolic shift, virulence is repressed and phenotypes associated with immune-evasion become established. Ultimately, copper acts not only as an antibacterial but as an environmental signal to drive differential metabolic and physiological changes to potentiate *S. aureus* adaptability and resilience to colonise multiple niches.

## Materials and Methods

### Bacterial strains and culture conditions

*S. aureus* mutants used in this study are derived from the USA300 JE2 background (Table 1)^59^. Transposon mutants from the Nebraska Transposon Library were confirmed by PCR and mutants were transduced into a clean USA300 JE2 background using phage Φ11. To construct a markerless *copL* mutant, the pTNT plasmid was transduced into copL:: *ΦNΣ* and allelic exchange was performed as described previously^30^. Strains were routinely cultured on Luria agar (LA) and in RPMI-A 1640 medium (Sigma-Aldrich; 0883) supplemented with 2% (v/v) RPMI amino acid solution (Sigma-Aldrich; R7131) (This is termed as RPMI-A). Cultures were grown at 37°C in 5% (v/v) CO_2_ under static conditions in 50mL Falcon tubes. For experimental assays, overnight cultures were centrifuged and resuspended in fresh RPMI-A to an optical density (OD600) of 0.1. When indicated, CuCl_2_ was added at the concentrations specified for each experiment. Erythromycin (10μg/mL; Sigma-Aldrich; E6376) was added where appropriate. Growth was monitored by measuring OD600 using either a UV/Vis spectrophotometer (Jenway) or a FLUOstar plate reader (BMG Labtech).

**Table 1:** Strains used in this study.

| Strain | Genotype | Reference |
| --- | --- | --- |
| **USA300 JE2 WT** | USA300 CA-MRSA isolate LAC without plasmids | Fey *et al.*, 2013 |
| **USA300 JE2 *copL*** | JE2 *copL::TNT* | This work, Bose *et al.*, 2013 |
| **USA300 JE2 *alsS*** | JE2 *alsS::ΦNΣ* | Bose *et al.*, 2013 |
| **USA300 JE2 *cidC*** | JE2 *cidC::ΦNΣ* | Bose *et al.*, 2013 |
| **USA300 JE2 *cidR*** | JE2 *cidR::ΦNΣ* | Bose *et al.*, 2013 |
| **USA300 JE2 *sigB*** | JE2 *sigB::ΦNΣ* | Bose *et al.*, 2013 |
| **USA300 JE2 *nirB*** | JE2 *nirB::ΦNΣ* | Bose *et al.*, 2013 |
| **USA300 JE2 *copL nirB*** | JE2 *copL::TNT nirB::ΦNΣ* | This work; Bose *et al.*, 2013 |

### Dissolved oxygen assay

Overnight cultures of *S. aureus* USA300 JE2 were diluted in fresh RPMI-A to OD600 of 0.1 in 10mL volumes and were incubated for 8 hours at 37°C under either static conditions in 5% CO_2_ or shaking conditions at 200rpm in ambient air. Independent cultures were prepared for each time point to avoid the disturbance of bacterial cultures during sampling. Dissolved oxygen concentrations were measured hourly using a ProODO optical dissolved oxygen meter (YSI/Xylem). To prevent atmospheric oxygen being introduced during the measurement and to avoid contamination of the sensor, culture samples were transferred into an anaerobic chamber (Don Whitley Scientific) and passed through a sterile 0.22μM syringe filter to remove bacterial cells. Dissolved oxygen was measured in the cell free supernatant and recorded as percentage dissolved oxygen. Uninoculated RPMI-A incubated under identical conditions served as controls. An anaerobic control was prepared by pre-equilibrating RPMI-A in an anaerobic chamber for 24 hours prior to measurement. Bacterial growth was monitored in parallel by OD600.

### Total RNA extraction

Overnight cultures of *S. aureus* WT and *copL* strains were grown in RPMI-A for 16 hours at 37 °C in 5 % CO_2_, then subcultured into fresh pre-warmed RPMI-A and grown to mid-exponential phase in the presence or absence CuCl_2_. RNA was stabilised using RNAprotect Bacteria Reagent (Qiagen; 76104) according to the manufacturer’s instructions and were stored at −80°C. Cells were lysed in 200µL TE buffer (10 mM TRIS-HCL pH 7.5, 10 mM EDTA) containing lysostaphin (10mg/mL; Sigma-Aldrich; L7386) for 30 min at 37 °C, followed by 10 µL of proteinase K treatment (10 mg/mL; Sigma-Aldrich; 1245680500) for 10 minutes. Lysates were transferred to Lysing Matrix B tubes (MP Biomedicals; 1169110-CF) and were homogenised in 600 µL of TRIzol reagent (Invitrogen; 15596026), using a FastPrep Instrument at 5m/s for 20 seconds. RNA was purified using an RNeasy Mini Kit (Qiagen; 74106), including an on-column DNase digestion and a subsequent Turbo DNA-free treatment (Invitrogen; AM1907).

### RNAseq and identification of differentially expressed genes

RNA integrity was assessed using an Agilent 2100 Bioanalyzer, and samples with RNA integrity number (RIN) ≥ 9 were used for sequencing. The RNA-seq libraries construction and sequencing were performed by the Oxford Genomics Centre using an Illumina NovaSeq 6000 sequencing at a depth of 20 million reads per sample following ribodepletion. Raw reads were checked for quality using FastQC. Adapter trimming was performed using Trimmomatic (v 0.36). Reads were aligned to the *S. aureus* USA300 FPR3757 reference genome (accession no. NC_007793.1) using HISAT2 (v 2.1.0) and the transcriptome was assembled using StringTie (v 1.3.3b). Differential gene expression analysis was conducted using DESeq2 in R. Gene expression changes are reported as log_2_ fold change relative to growth in RPMI-A without copper unless otherwise stated. Genes with a log_2_ fold change of > 1 or < −1 with an adjusted p-value ≤ 0.01 were considered significantly differentially expressed.

### Gene set enrichment analysis (GSEA)

Gene set enrichment analysis (GSEA v4.2.3) was performed on the gene count matrix and formatted into a GCT file with corresponding phenotype labels in CLS format. Custom gene sets were constructed using KEGG-derived functional categories for *S. aureus* USA300 and independently defined I-modulons established previously^32^. Orthologue mapping was performed using aureowiki as a key source (https://aureowiki.med.uni-greifswald.de/Main_Page)^60^. The created gene sets were compiled into a GMX format and annotated using CHIP annotation files. Enrichment results were filtered to include the gene sets with a false discovery rate (FDR) q-value of <0.05.

### Quantitative RT-PCR

Complementary DNA (cDNA) was synthesised from 1000ng total RNA using the Superscript IV VILO reagent kit (Invitrogen; 11756050). Quantitative PCR reactions contained 5μL SYBR Green Master (Mix Applied Biosystems; 4385612), 300nM primer and 1ng of cDNA. Primer pairs are shown (Table 2). Reactions were done using the QuantStudio 3 Real-Time PCR System (Applied Biosystems; A28567) in a MicroAmp optical 96-well plate (Applied Biosystems; N8010560) and sealed with the optical adhesive film (Applied Biosystems; 4311971). PCR reactions were done with standard cycling conditions (95 °C denaturation and 60 °C annealing). Relative gene expression was calculated using the 2^-ΔΔct^ method with *gyrB* as the reference gene^61^.

**Table 2:**
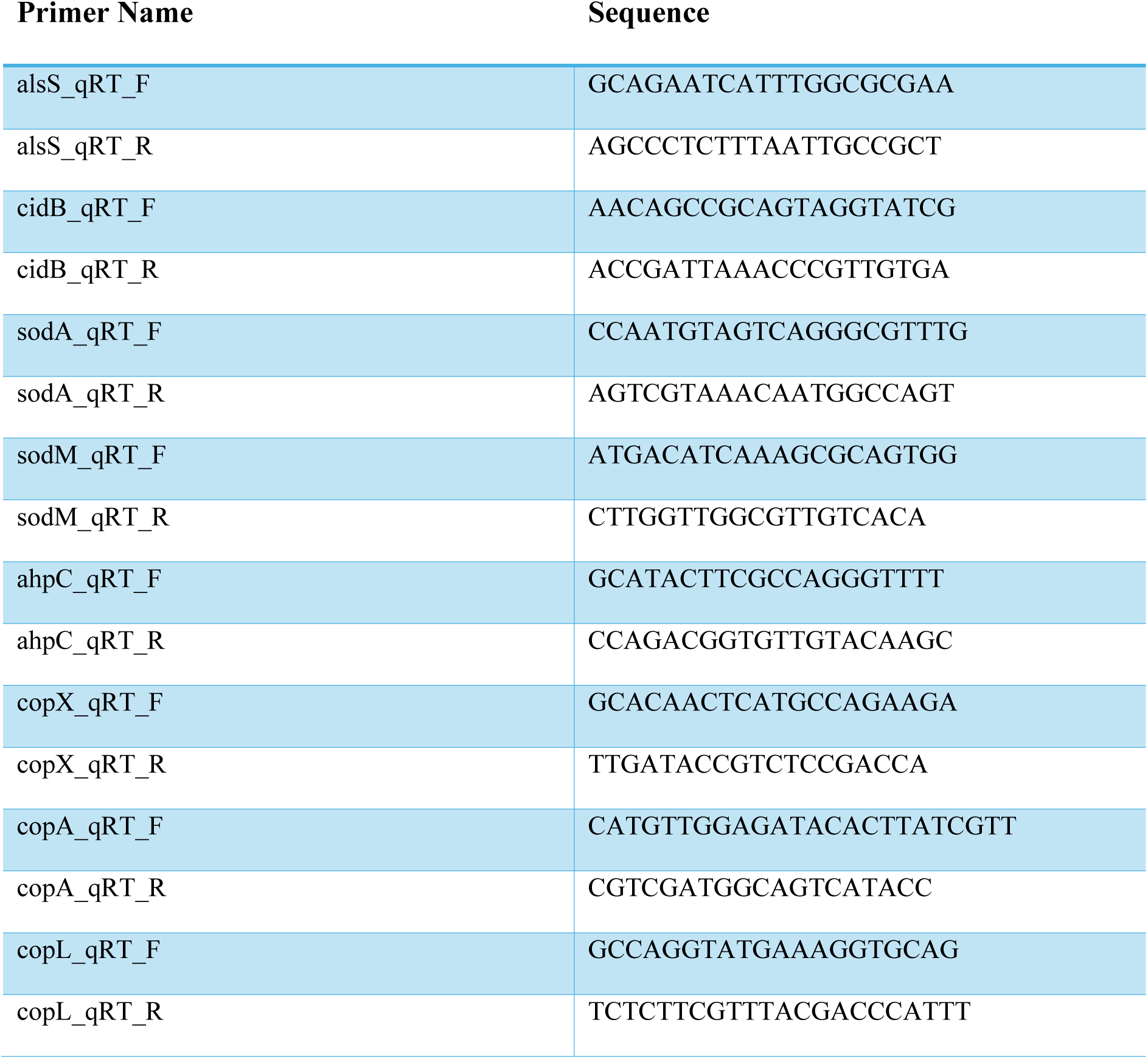
qRT-PCR primers used in this study.

### Determination of intracellular pH

Overnight cultures of *S. aureus* strains were prepared in 10 mL of RPMI-A (including appropriate antibiotics) and incubated at 37 °C + 5 % CO_2_ for 16 hours. Cultures were pelleted and resuspended in sterile cold 10 mM phosphate buffered saline containing, 1.5 g/L Na_2_HPO_4_ (Merck; 567550), 0.22 g/L of NaH_2_PO_4_ (Merck; S0751), and 8.5 g/L of NaCl (Merck; S9888) and adjusted to a final pH of 7.4 and to a final OD600 of 0.6. The cells were stained with 10µM of carboxyfluorescein diacetate succinimidyl ester (cFDA-SE) (Sigma-Aldrich; 21888) and incubated at 37 °C for 30 min. The cells were centrifuged at 4000 rpm followed and resuspended in 50 mM potassium phosphate buffer (pH 6.0) containing 10 mM glucose. The cells were then incubated at 30 °C for 30 min. Cells were centrifuged and resuspended in 50 mM potassium phosphate buffer containing 10 mM glucose and kept on ice until required. Stained cells were resuspended in RPMI-A to an OD600nm of 0.1. The samples were incubated statically at 37 °C + 5% CO_2_ for 6 hours in a 96-well plate with either 100μM or 500μM CuCl_2_ at a final volume of 200μL. The fluorescence emission ratios (490nm/435nm) were measured using a Hidex Sense Plate Reader (Hidex). The intracellular pH was determined using calibration curves generated from stained cells permeabilised with ethanol (63% v/v) for 30 min. Cells were pelleted and resuspended in RPMI-A adjusted for pH ranging from 3.5 – 8.5 (in 0.5 increments), using HCL or NaOH.

### Measurement of end metabolites

Mid-exponentially growing *S. aureus* in +/- 100 μM CuCl_2_ were pelleted by centrifugation, and the supernatants were used to determine the extracellular concentration of metabolites. Extracellular acetate and pyruvate concentrations were determined using the colorimetric acetate (Sigma-Aldrich; MAK086) and pyruvate kits (Abcam; AB65342) according to the manufacturer’s instructions. Acetoin concentrations were determined using a modified quantitative colorimetric acetoin assay as described previously^62^. A standard curve ranging from 0 – 150 µM acetoin was prepared in RPMI-A from a 1 mM stock concentration of pure acetoin (Sigma-Aldrich; W200832) dissolved in distilled H_2_O. To 200 µL of sample 140 µL creatine (0.5 % w/v in H_2_O) (Sigma-Aldrich; C3630), 200 µL α-naphthol (5 % w/v in 95 % (v/v) ethanol) (Sigma-Aldrich; N1000) and 200 µL potassium hydroxide (KOH, 40 % w/v in H_2_O) were added sequentially. The samples were then vortexed and incubated at room temperature for 15 mins. The samples were then vortexed again and the OD 560nm was measured using a FLUOstar Omega plate reader at a final volume of 200µL. Extracellular total Lactate and Glucose concentrations were determined using a lactic acid (Megazyme; K-DLATE), and glucose (Megazyme; K-GLUC) assay kits following manufacturer’s instructions.

To determine the intracellular concentration of succinate, following centrifugation of mid exponentially growing cells, the supernatant was removed, and the pellets were washed twice in PBS before being resuspended in 500 µL succinate assay buffer provided in the colorimetric assay kit (Sigma-Aldrich; MAK184). The samples were incubated at 80 °C for 20 minutes, transferred to a Lysing Matrix B tube and homogenised using the MP Biomedical Fast Prep Instrument at a frequency of 5M/second for 20 seconds followed by a further homogenisation of 4M/second for 20 seconds. Samples were transferred to a fresh microcentrifuge and centrifuged at 13,000 x g for 10 minutes. Supernatants were transferred to a fresh tube and kept on ice until use. The samples were diluted 1:2 in succinate assay buffer and the succinate concentrations were determined according to the manufacturer’s instructions.

For measuring the concentrations of glutamate and glutamine, *S. aureus* was grown to mid-exponential phase and then washed twice in PBS. After washing the bacterial cells were diluted 1:20 in PBS at a final volume of 1mL and the samples were then assayed as described in the Glutamine/Glutamate-Glo Assay (Promega; J8021) standard protocols. Luminescence was measured with the FLUOstar Omega plate reader using white, clear bottomed 96 well assay plate (Corning; CLS3610).

### Supplement dose-response growth curves

Overnight cultures of *S. aureus* WT and *copL* mutants were subcultured in fresh RPMI-A to an OD600 of 0.1. CuCl_2_ was then added to the bacterial culture to achieve a final concentration of 2mM. For supplementation, D- and L- glutamic acid (Sigma-Aldrich; G1001and G1251) as well as D- and L- proline (Sigma-Aldrich; 858919 and P0380) were prepared fresh in RPMI-A prior to each experiment. Solutions were filter sterilised using a 0.22μM sterile filter. When required, amino acids were fully dissolved by vortexing prior to filtration. Serial dilutions of each supplement were prepared directly in a flat-bottom 96-well microtiter plate to a final volume of 100μL. Bacterial cultures with or without added copper were then added at a volume of 100μL to each well to a final volume of 200μL and a final copper concentration of 1mM. Plates were sealed with a gas-permeable Breathe-Easy membrane (Sigma-Aldrich; Z380059) and incubated at 37°C in 5% CO_2._ OD600 was measured every 30 minutes for 24 hours using a FLUOstar plate reader. Growth analysis was performed in R studio.

### Nitrite concentration assay

For nitrite concentration, the nitrate reduction test was used (Sigma-Aldrich; 73426) and modified as follows. A standard curve of sodium nitrite (Sigma-Aldrich; 237213) ranging from 1 - 200 μM was prepared in ddH2O to serve as calibration standards. A 1:1 ratio of sulfanilic acid and N,N-Dimethyl-1-1naphthylamine from the kit were mixed and used fresh for the experiment, 20uL of this Greiss mix was added to 230uL of ddH2O and 30uL of sample supernatant or standard in a 96 well plate. Samples were incubated for 30 minutes, then optical density was measured at 548nm. The nitrite concentration of the sample was then determined from the calibration standard curve.

### Statistical analyses

All statistical analyses of data were conducted using the Graphpad Prism 9 software and results with a *P-*value ≤ 0.05 were considered significant. T-tests were used when comparing two groups. A One-way ANOVA was used when comparing 3 or more groups of a single variable. A two-way ANOVA was conducted when comparing groups of two variables. Details of tests and post-tests (Dunnet’s, Tukey and Sidak) used are given in the figure legends.

## Data Availability

All data supporting the findings of this study are publicly available in open repositories. Raw RNA sequencing data have been deposited in the NCBI Gene Expression Omnibus (GEO) under accession number GSE335600.

All numerical data underlying the figures and statistical analyses are available on the University of Leicester Figshare repository (https://doi.org/10.25392/leicester.data.32687358). This repository includes all processed datasets used to generate figures. These files enable reproduction of the reported statistical analyses and graphical outputs.

## Acknowledgements

SB was funded by NIHR Health Protection Research Unit in Chemical Threats and Hazards (Award No. NIHR207293), a partnership with the Health and Safety Executive and the UK Health Security Agency, also funding from the NIHR Leicester Biomedical Research Centre (BRC). The views expressed are those of the authors and not necessarily those of the NHS, the NIHR, the Department of Health and Social Care, the Health and Safety Executive or the UK Health Security Agency. IK was funded by a BBSRC Midlands Integrative Biosciences Training Partnership PhD studentship. JP, DS were funded by BBSRC Grant BB/S006818/1. We thank Professor Mick Whelan and Dr Helen Crabb for support of SB during his PhD studentship. For the purpose of open access, the author has applied a Creative Commons Attribution license (CC BY) to any Author Accepted Manuscript version arising from this submission.

## Author contribution

SCB, IK, JP, DS, HRS: Investigation, Formal Analysis, Methodology, Data Curation, Visualization, Writing – Original Draft; JG, JMK, PWA, KW, JAM: Conceptualization, Visualization, Formal Analysis, Funding Acquisition Supervision, Writing – Review & Editing.

## Ethical approval, permission to reproduce material, patient consent

not required.

