## Supplementary file for "Microaerobic Copper Stress Redirects Pyruvate Metabolism and Reveals a CopL-linked Nitrogen Response in *Staphylococcus aureus*"

**A**

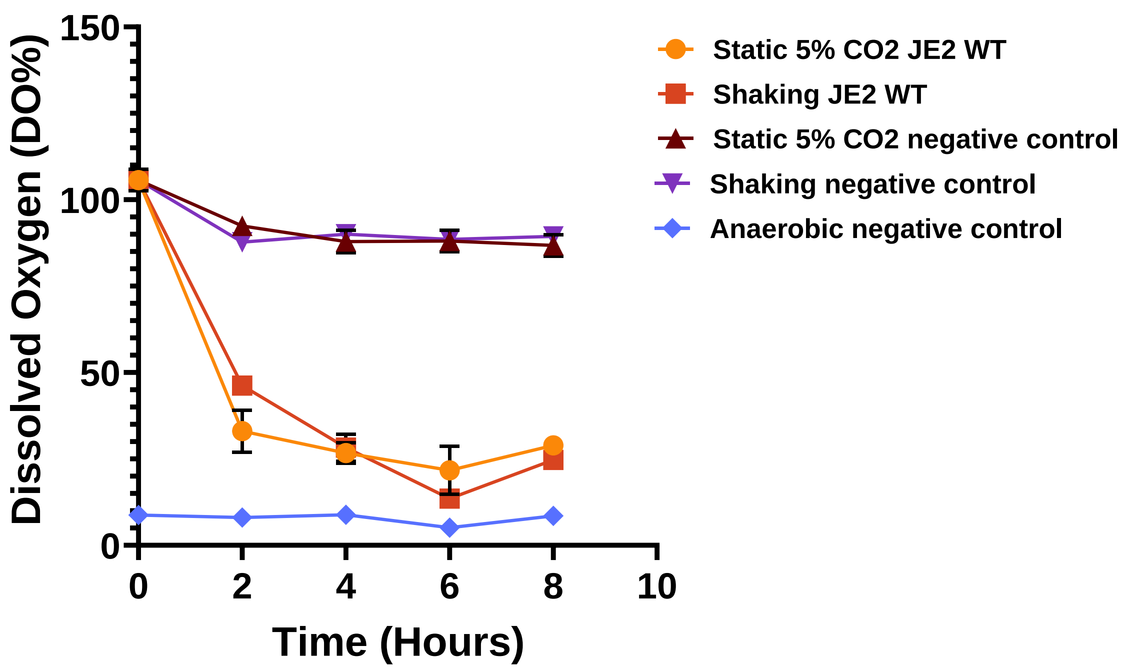


**B**

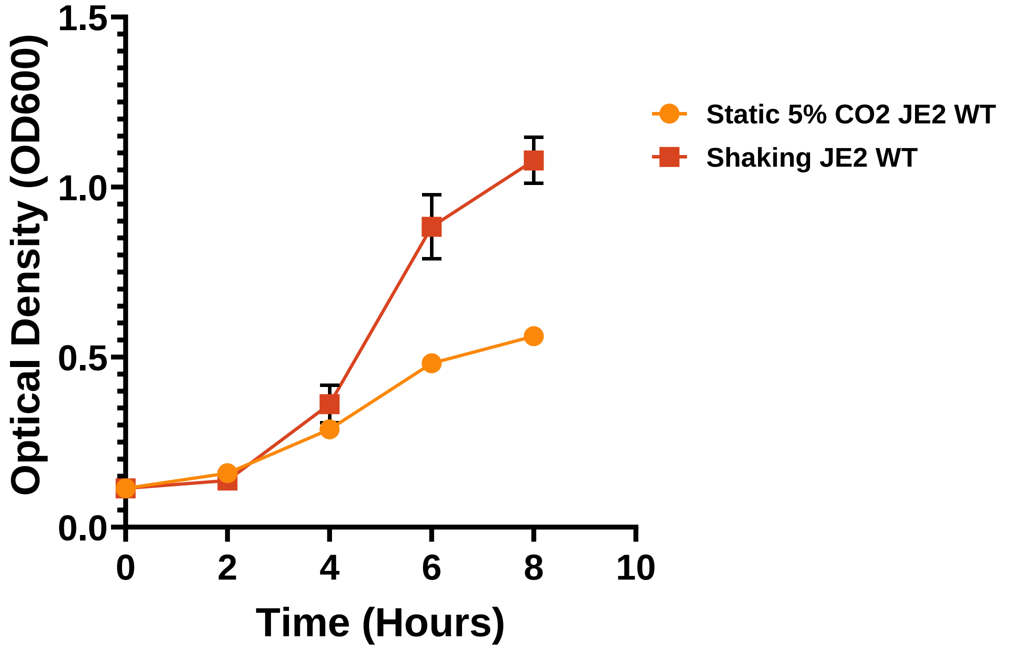


**Supplementary figure 1: Growth of *S. aureus* reduces dissolved oxygen, with slower growth under static microaerobic (5% $CO_2$) conditions.**

(A) Dissolved oxygen levels were measured in filter bacterial cultures using a ProODO oxygen probe. Sterile medial controls were included to validate probe performance. Error bars represent the ± standard error of the mean (SEM) from at least 3 biological replicates.

(B) Growth curve of WT *S. aureus* under static 5% $CO_2$ and aerobic shaking conditions (160 RPM). Absorbance was measured at OD600. Error bars represent the ± standard error of the mean (SEM) from at least 3 biological replicates.

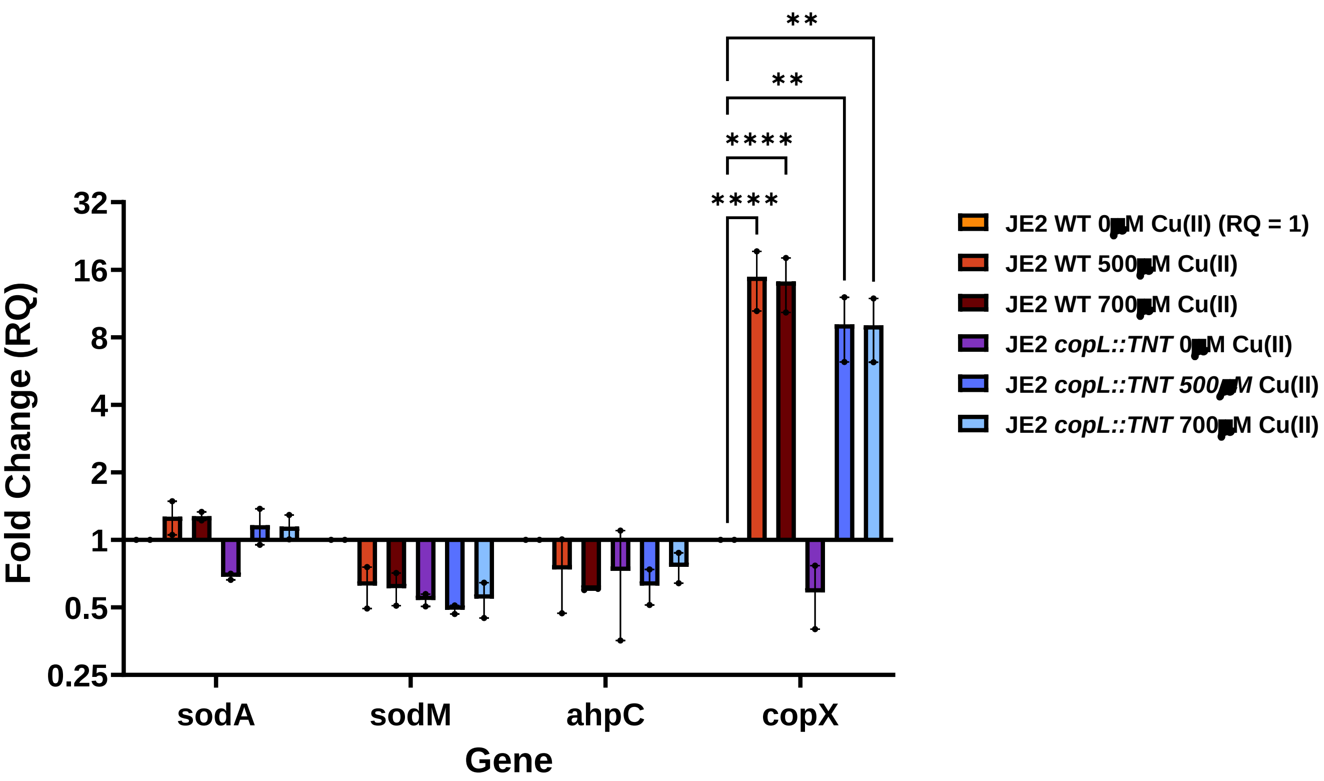


**Supplementary figure 2: Expression of genes related to oxidative stress in response to copper shock in WT and *copL* strain**

Transcription of genes *sodA*, *sodM*, *ahpC* related to oxidative stress were investigated in the WT and *copL* strain when shocked with either 500 or 700 µM CuCl2. Expression of *copX* was investigated as a positive control to ensure a copper response would be achieved under these conditions. Results were determined by qRT-PCR of RNA extracted from cells cultured at 37 °C + 5 % CO2 in RPMIA to mid exponential phase and shocked for 10-minutes with either 500 or 700µM $CuCl_2$.Relative expression was calculated as RQ using the ΔΔCt method, which normalises expression of each strain against an endogenous control (*gyrB*) and expresses the data relative to a reference strain (JE2 WT in the absence of copper; RQ = 1). Error bars represent the ± standard error of mean (SEM) from two biological replicates. Significance between conditions was determined using a two-way ANOVA with Dunnett's multiple comparison test (ns = no significant difference, ** P < 0.01, **** P < 0.001).

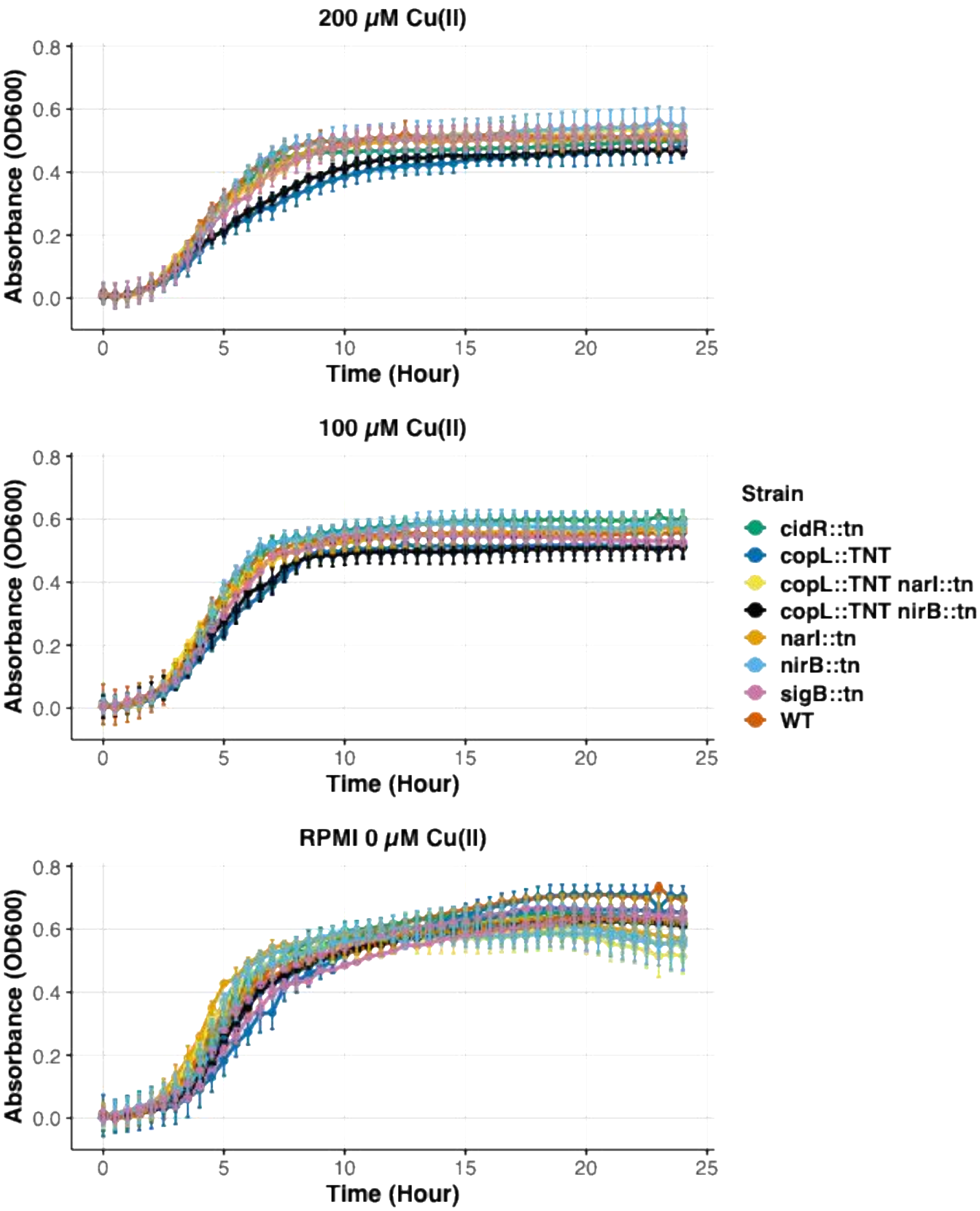


**Supplementary figure 3: The effect of $CuCl_2$ on JE2 mutants in RPMI-A under 5% $CO_2$ to determine subinhibitory concentrations**

*S. aureus* JE2 WT and relevant regulatory and metabolic mutants were cultured in RPMI-A with either either 0µM, 100µM or 200µM Cu(II)$Cl_2$. Growth curves were conducted in flat bottomed 96-well plates with a Breathe-Easy polyurethane membrane and incubated in an Omega plate reader with the atmosphere set at 5% CO2 atmosphere at 37°C. OD600 measurements were taken every 30 minutes. Error bars represent ± standard error of mean (SEM) from 2 independent biological replicates.

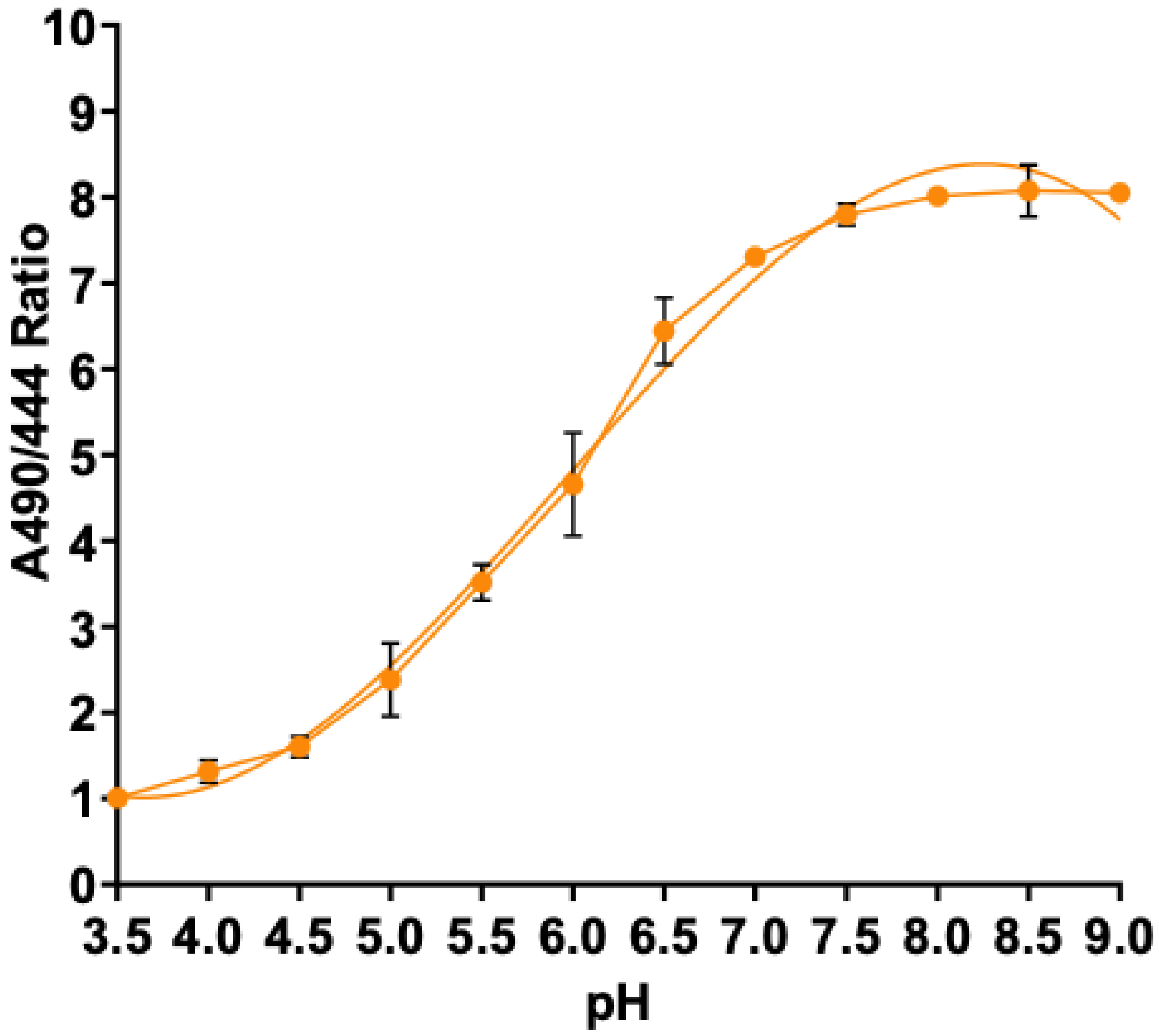


**Supplementary figure 4: Determining the standard curve to measure the Intracellular pH of *S. aureus* USA300 WT and *copL* mutant grown in the presence and absence of copper.**

The A490/444 nm ratio of fluorescence of *S. aureus* USA300 WT, stained with CFDA-SE, permeabilised in ethanol, and resuspended in media of various pH, was measured and plotted. Error bars represent the ± standard error of the mean (SEM) from 3 independent biological replicates.

A

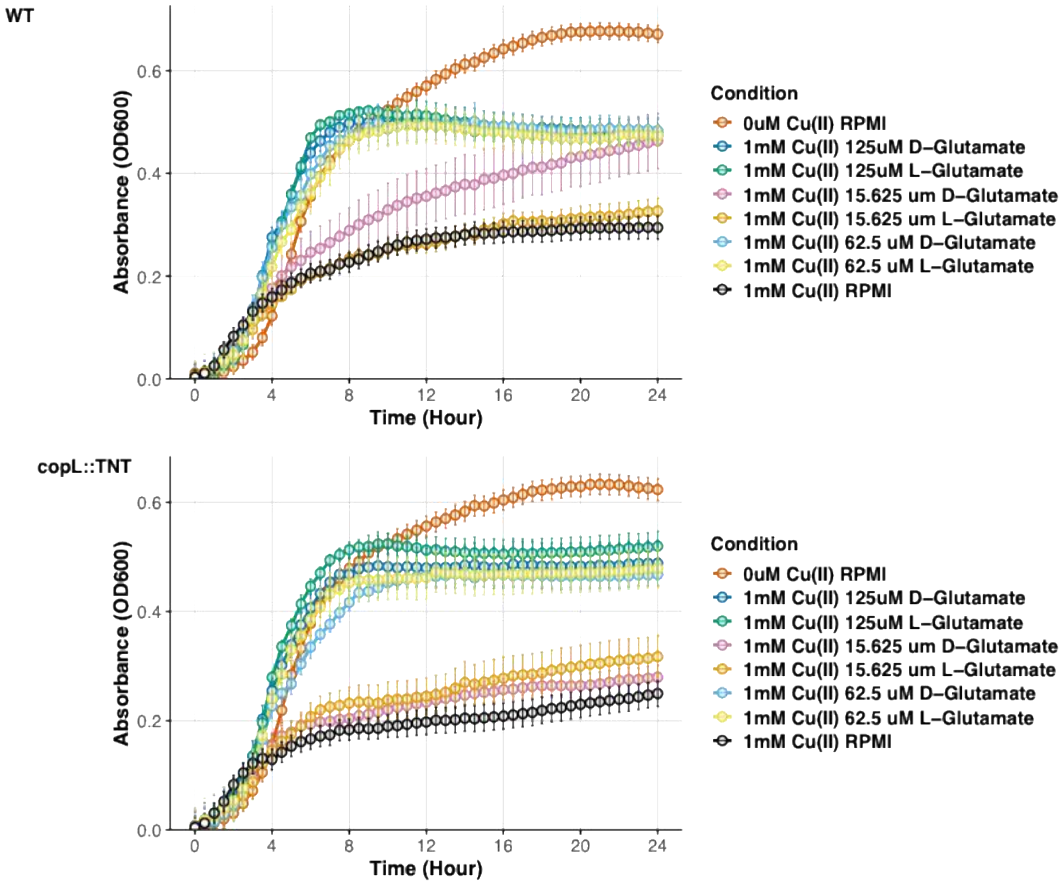


B

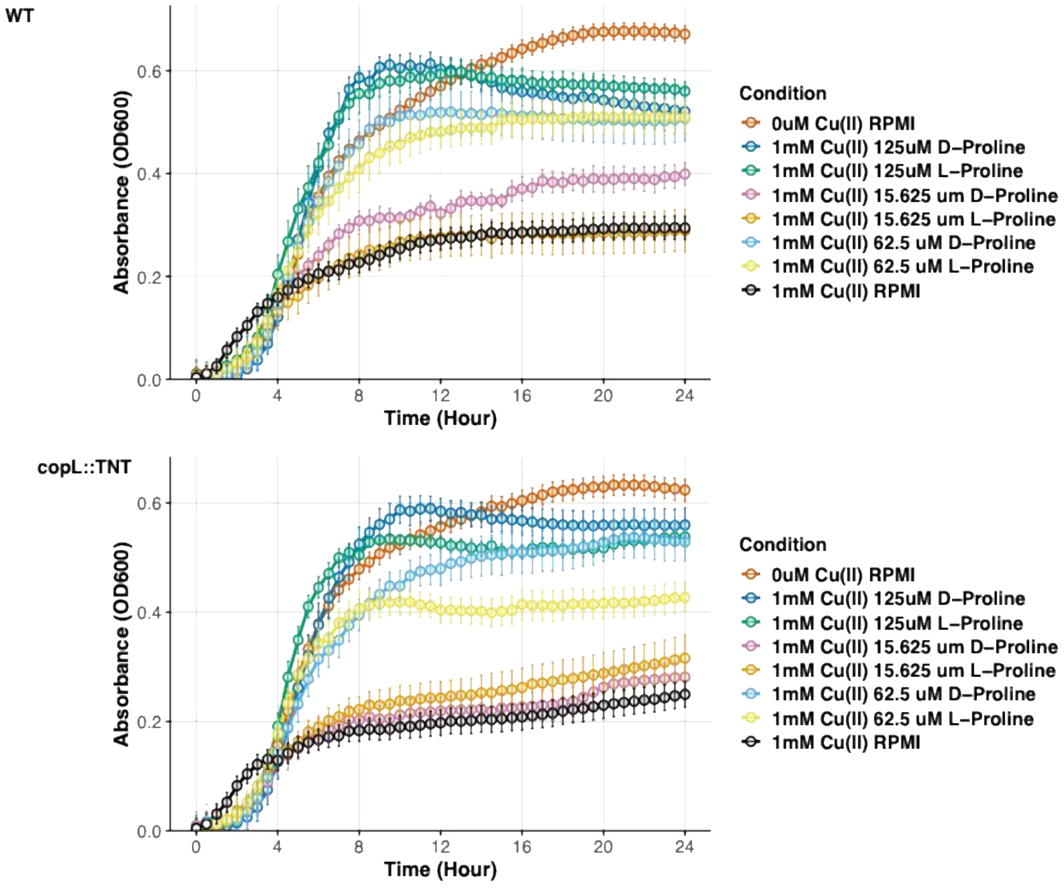


**Supplementary figure 5: Glutamate and proline are protective of staphylococcal growth against toxic copper concentrations.**

*S. aureus* JE2 WT and *copL* mutant were cultured in RPMI-A supplemented with 1mM toxic Cu(II)$Cl_2$ concentrations and either (A) D-glutamate or L-glutamate, (B) D-proline or L-proline at increasing concentrations. The total concentrations ranged from 125µM -15.625µM for both amino acids. Growth curves were conducted in flat bottomed 96-well plates with a Breathe-Easy polyurethane membrane and incubated in an Omega plate reader with the atmosphere set at 5% CO2 atmosphere at 37°C. OD600 measurements were taken every 30 minutes. Error bars represent ± standard error of mean (SEM) from 3 independent biological replicates.

*Table 1* The differentially expressed genes of *Staphylococcus aureus* in response 100μM $CuCl_2$ under microaerobic conditions *(adjusted $p \leq 0.01$; $log_2$ fold change $\geq 1$ or $\leq -1$)*. The locus tag and the corresponsing Gene name and Protein product are displayed alongside the biological function.

| LOCUS TAG | GENE NAME | PRODUCT DESCRIPTION | BIOLOGICAL FUNCTION | LOG FOLD CHANGE | ADJUSTED P-VALUE |
|---|---|---|---|---|---|
| **SAUSA300_RS13750** | cidB | LrgB family protein | Cell lysis/autolysis | 1.73 | 1.99E-76 |
| **SAUSA300_RS13755** | cidA | holin-like murein hydrolase modulator | Cell lysis/autolysis | 1.73 | 1.99E-76 |
| **SAUSA300_RS00395** | copX | copper-translocating P-type ATPase | Copper homeostasis | 4.42 | 5.61E-289 |
| **SAUSA300_RS00400** | copL | DUF1541 domain-containing protein | Copper homeostasis | 4.23 | 6.75E-204 |
| **SAUSA300_RS13855** | copA | copper-translocating P-type ATPase | Copper homeostasis | 3.84 | 0.00E+00 |
| **SAUSA300_RS13860** | copZ | copper chaperone | Copper homeostasis | 3.79 | 2.95E-278 |
| **SAUSA300_RS13845** | x | maltose O-acetyltransferase | Metabolism | 1.08 | 8.52E-21 |
| **SAUSA300_RS13375** | x | SDR family oxidoreductase | Metabolism | 1.13 | 1.32E-09 |
| **SAUSA300_RS12320** | fdhD | formate dehydrogenase accessory sulfurtransferase | Metabolism | 1.00198846 | 4.69E-11 |
| **SAUSA300_RS02320** | cysM/mccA | cysteine synthase family protein | Metabolism (Amino acid) | 1.81 | 4.25E-03 |
| **SAUSA300_RS02325** | metB/mccB | bifunctional cystathionine gamma-lyase/homocysteine desulfhydrase | Metabolism (Amino acid) | 1.81 | 4.25E-03 |
| **SAUSA300_RS00995** | idpC | alpha-keto acid decarboxylase family protein | Metabolism (Amino acid) | 1.58 | 1.75E-07 |
| **SAUSA300_RS04665** | argG | argininosuccinate synthase | Metabolism (Amino acid) | 1.22 | 6.74E-15 |
| **SAUSA300_RS04660** | argH | argininosuccinate lyase | Metabolism (Amino acid) | 1.22 | 6.74E-15 |
| **SAUSA300_RS02985** | vraA | long-chain fatty acid--CoA ligase | Metabolism (Fatty acid) | 2.14 | 3.31E-38 |
| **SAUSA300_RS03815** | bmrU | lipid kinase | Metabolism (Lipid) | 1.07 | 4.33E-12 |
| **SAUSA300_RS11940** | alsS | acetolactate synthase | Metabolism (Pyruvate) | 2.87 | 5.32E-39 |

| | | | | | |
|---|---|---|---|---|---|
| **SAUSA300_RS11935** | budA/alsD | acetolactate decarboxylase | Metabolism (Pyruvate) | 2.56 | 3.40E-38 |
| **SAUSA300_RS13745** | cidC | pyruvate oxidase | Metabolism (Pyruvate) | 1.40 | 5.95E-76 |
| **SAUSA300_RS06765** | acnA/citB | aconitate hydratase | Metabolism (TCA cycle) | 1.03 | 1.48E-10 |
| **SAUSA300_RS11675** | tRNA-Glu | tRNA-Glutamate | Translation | 1.30 | 2.19E-07 |
| **SAUSA300_RS11680** | tRNA-Asn | tRNA-Aspartate | Translation | 1.25 | 1.16E-06 |
| **SAUSA300_RS11670** | tRNA-Val | tRNA-Valine | Translation | 1.04 | 2.98E-07 |
| **SAUSA300_RS11660** | tRNA-Gln | tRNA-Glutamine | Translation | 1.01 | 3.56E-06 |
| **SAUSA300_RS11655** | tRNA-Lys | tRNA-Lysine | Translation | 1.00 | 1.86E-05 |
| **SAUSA300_RS12940** | cobA | uroporphyrinogen-III C-methyltransferase | Nitrogen metabolism | 1.74 | 5.49E-03 |
| **SAUSA300_RS12945** | nirD | nitrite reductase small subunit | Nitrogen metabolism | 1.74 | 5.49E-03 |
| **SAUSA300_RS04245** | ohr | organic hydroperoxide resistance protein | Oxidative stress | 1.02 | 6.22E-16 |
| **SAUSA300_RS04510** | dltX | teichoic acid D-Ala incorporation-associated protein | Resistance to cationic antimicrobial peptides | 2.03 | 1.99E-91 |
| **SAUSA300_RS04515** | dltA | D-alanine--poly(phosphoribitol) ligase subunit | Resistance to cationic antimicrobial peptides | 1.76 | 6.08E-56 |
| **SAUSA300_RS04520** | dltB | PG:teichoic acid D-alanyltransferase | Resistance to cationic antimicrobial peptides | 1.76 | 6.08E-56 |
| **SAUSA300_RS04525** | dltC | D-alanine--poly(phosphoribitol) ligase subunit 2 | Resistance to cationic antimicrobial peptides | 1.44 | 1.01E-28 |
| **SAUSA300_RS04530** | dltD | D-alanyl-lipoteichoic acid biosynthesis protein | Resistance to cationic antimicrobial peptides | 1.44 | 1.01E-28 |
| **SAUSA300_RS03470** | vraF | ABC transporter ATP-binding protein | Resistance to cationic antimicrobial peptides | 1.16 | 8.59E-64 |
| **SAUSA300_RS03475** | vraG | FtsX-like permease family protein | Resistance to cationic antimicrobial peptides | 1.16 | 8.59E-64 |
| **SAUSA300_RS06820** | mprF | bifunctional lysylphosphatidylglycerol flippase/synthetase | Resistance to cationic antimicrobial peptides | 1.03 | 1.87E-19 |
| **SAUSA300_RS11820** | opuD2 | BCCT family transporter | Transport | 1.31 | 2.01E-12 |

| | | | | | |
|---|---|---|---|---|---|
| **SAUSA300_RS00915** | ssuB | ABC transporter ATP-binding protein | Transport | 1.14 | 1.22E-03 |
| **SAUSA300_RS06965** | pstS | phosphate ABC transporter substrate-binding protein | Transport (Phosphate) | 1.33 | 9.81E-06 |
| **SAUSA300_RS03005** | vraX | protein VraX | Virulence | 4.02 | 1.46E-121 |
| **SAUSA300_RS05670** | ecb | complement convertase inhibitor Ecb | Virulence | 2.32 | 0.0023 |
| **SAUSA300_RS05690** | efb | fibrinogen-binding protein | Virulence | 1.84 | 2.73E-06 |
| **SAUSA300_RS05695** | scc | complement inhibitor SCIN-B | Virulence | 1.56 | 1.68E-07 |
| **SAUSA300_RS00990** | entB | isochorismatase family protein | Virulence | 1.42 | 1.38E-07 |
| **SAUSA300_RS13070** | hlgA | bi-component gamma-hemolysin HlgAB subunit A | Virulence | 1.12 | 4.03E-04 |
| **SAUSA300_RS10620** | X | phage head-tail adapter protein | Phage SA3 | 1.12 | 8.13E-17 |
| **SAUSA300_RS10625** | X | phage head-tail adapter protein | Phage SA3 | 1.12 | 8.13E-17 |
| **SAUSA300_RS10605** | X | phage tail protein | Phage SA3 | 1.12 | 5.25E-09 |
| **SAUSA300_RS10635** | X | phage major capsid protein | Phage SA3 | 1.03330884 | 1.08E-11 |

| LOCUS TAG | GENE NAME | PRODUCT DESCRIPTION | BIOLOGICAL FUNCTION | LOG FOLD CHANGE | ADJUSTED P-VALUE |
|---|---|---|---|---|---|
| **SAUSA300_RS04580** | X | SidA/IucD/PvdA family monooxygenase | DNA stress | -3.3783203 | 3.00E-12 |
| **SAUSA300_RS02020** | ahpF | alkyl hydroperoxide reductase subunit F | DNA stress | -1.16 | 9.23E-11 |
| **SAUSA300_RS09410** | comK | competence protein | DNA stress | -1.24 | 2.39E-10 |
| **SAUSA300_RS04985** | comK1 | competence protein ComK | DNA stress | -2.31 | 2.87E-07 |
| **SAUSA300_RS06710** | lexA | transcriptional repressor | DNA stress | -1.14 | 3.67E-09 |
| **SAUSA300_RS09405** | sigS | RNA polymerase sigma factor | DNA stress | -1.18 | 6.24E-09 |
| **SAUSA300_RS06840** | umuC | Y-family DNA polymerase | DNA stress | -1.64 | 1.12E-09 |
| **SAUSA300_RS00760** | phnD | phosphonate ABC transporter substrate-binding protein | Metabolism | -1.17 | 6.12E-06 |
| **SAUSA300_RS13365** | cntK | histidine racemase | Metabolism (Amino acid) | -1.40 | 4.51E-15 |
| **SAUSA300_RS14515** | hisD | histidinol dehydrogenase | Metabolism (Amino acid) | -1.27 | 5.57E-06 |

| | | | | | |
|---|---|---|---|---|---|
| **SAUSA300_RS14520** | hisG | ATP phosphoribosyltransferase | Metabolism (Amino acid) | -1.27 | 5.57E-06 |
| **SAUSA300_RS14525** | hisZ | ATP phosphoribosyltransferase regulatory subunit | Metabolism (Amino acid) | -1.33 | 3.53E-05 |
| **SAUSA300_RS11075** | ilvA | threonine ammonia-lyase | Metabolism (Amino acid) | -1.20 | 1.05E-10 |
| **SAUSA300_RS11040** | ilvB | biosynthetic-type acetolactate synthase large subunit | Metabolism (Amino acid) | -1.34 | 2.95E-06 |
| **SAUSA300_RS11050** | ilvC | ketol-acid reductoisomerase | Metabolism (Amino acid) | -1.35 | 2.01E-06 |
| **SAUSA300_RS11035** | ilvD | dihydroxy-acid dehydratase | Metabolism (Amino acid) | -1.44 | 2.40E-06 |
| **SAUSA300_RS11045** | ilvH | ACT domain-containing protein | Metabolism (Amino acid) | -1.34 | 2.95E-06 |
| **SAUSA300_RS11055** | leuA | 2-isopropylmalate synthase | Metabolism (Amino acid) | -1.37 | 5.71E-08 |
| **SAUSA300_RS11060** | leuB | 3-isopropylmalate dehydrogenase | Metabolism (Amino acid) | -1.17 | 9.78E-10 |
| **SAUSA300_RS11065** | leuC | 3-isopropylmalate dehydratase large subunit | Metabolism (Amino acid) | -1.28 | 2.20E-11 |
| **SAUSA300_RS11070** | leuD | 3-isopropylmalate dehydratase small subunit | Metabolism (Amino acid) | -1.22 | 1.79E-09 |
| **SAUSA300_RS01905** | metC | PLP-dependent transferase | Metabolism (Amino acid) | -1.35 | 1.34E-10 |
| **SAUSA300_RS01895** | metE | 5-methyltetrahydropteroyltriglutamate--homocysteine S-methyltransferase | Metabolism (Amino acid) | -1.35 | 1.34E-10 |
| **SAUSA300_RS01900** | metF | bifunctional homocysteine S-methyltransferase/methylenetetrahydrofolate reductase | Metabolism (Amino acid) | -1.35 | 1.34E-10 |
| **SAUSA300_RS01910** | metI | aminotransferase class I/II-fold pyridoxal phosphate-dependent enzyme | Metabolism (Amino acid) | -1.35 | 1.34E-10 |
| **SAUSA300_RS04300** | metN2 | methionine ABC transporter ATP-binding protein | Metabolism (Amino acid) | -2.61 | 5.89E-31 |
| **SAUSA300_RS04305** | metP1 | ABC transporter permease | Metabolism (Amino acid) | -2.61 | 5.89E-31 |
| **SAUSA300_RS04310** | metQ1 | MetQ/NlpA family ABC transporter substrate-binding protein | Metabolism (Amino acid) | -2.53 | 8.90E-36 |
| **SAUSA300_RS00980** | rocD | ornithine--oxo-acid transaminase | Metabolism (Amino acid) | -1.13 | 7.59E-09 |
| **SAUSA300_RS06640** | thrD | aspartate kinase | Metabolism (Amino acid) | -1.07 | 7.55E-04 |
| **SAUSA300_RS06855** | trpE | anthranilate synthase component I | Metabolism (Amino acid) | -1.00 | 7.64E-07 |
| **SAUSA300_RS06860** | trpG | aminodeoxychorismate/anthranilate synthase component II | Metabolism (Amino acid) | -1.00 | 7.64E-07 |
| **SAUSA300_RS02025** | ahpC | peroxiredoxin | Oxidative stress | -1.22 | 7.03E-10 |
| **SAUSA300_RS04810** | opp4A | ABC transporter substrate-binding protein | Transport | -1.05 | 2.30E-03 |
| **SAUSA300_RS00060** | x | AzlD domain-containing protein | Transport | -1.15 | 7.38E-05 |

| | | | | | |
|---|---|---|---|---|---|
| **SAUSA300_RS00055** | x | AzlC family ABC transporter permease | Transport | -1.15 | 7.38E-05 |
| **SAUSA300_RS11525** | dps | DNA starvation/stationary phase protection protein | Transport (Iron) | -2.40 | 3.25E-03 |
| **SAUSA300_RS10250** | ftnA | H-type ferritin FtnA | Transport (Iron) | -2.07 | 3.98E-04 |
| **SAUSA300_RS05535** | isdB | heme uptake protein | Transport (Iron) | -1.44 | 2.94E-11 |
| **SAUSA300_RS05545** | isdC | heme uptake protein | Transport (Iron) | -1.04 | 5.11E-27 |
| **SAUSA300_RS05550** | isdD | hypothetical protein | Transport (Iron) | -1.04 | 5.11E-27 |
| **SAUSA300_RS05555** | isdE | heme ABC transporter substrate-binding protein | Transport (Iron) | -1.04 | 5.11E-27 |
| **SAUSA300_RS05560** | isdF | iron ABC transporter permease | Transport (Iron) | -1.18 | 1.44E-36 |
| **SAUSA300_RS05570** | isdG | staphylobilin-forming heme oxygenase | Transport (Iron) | -1.03 | 5.99E-24 |
| **SAUSA300_RS00610** | sbnA | 2,3-diaminopropionate biosynthesis protein | Transport (Iron) | -1.35 | 1.45E-15 |
| **SAUSA300_RS00615** | sbnB | N-[(2S)-2-amino-2-carboxyethyl]-L-glutamate dehydrogenase | Transport (Iron) | -1.35 | 1.45E-15 |
| **SAUSA300_RS00620** | sbnC | staphyloferrin B biosynthesis protein | Transport (Iron) | -1.18 | 5.56E-12 |
| **SAUSA300_RS00625** | sbnD | staphyloferrin B export MFS transporter | Transport (Iron) | -1.18 | 5.56E-12 |
| **SAUSA300_RS00630** | sbnE | L-2,3-diaminopropanoate--citrate ligase | Transport (Iron) | -1.18 | 5.56E-12 |
| **SAUSA300_RS00635** | sbnF | 3-(L-alanin-3-ylcarbamoyl)-2-[(2-aminoethylcarbamoyl)methyl]-2-hydroxypropanoate synthase | Transport (Iron) | -1.18 | 5.56E-12 |
| **SAUSA300_RS00640** | sbnG | staphyloferrin B biosynthesis citrate synthase | Transport (Iron) | -1.18 | 5.56E-12 |
| **SAUSA300_RS00645** | sbnH | staphyloferrin B biosynthesis decarboxylase | Transport (Iron) | -1.18 | 5.56E-12 |
| **SAUSA300_RS00650** | sbnI | bifunctional transcriptional regulator/O-phospho-L-serine synthase | Transport (Iron) | -1.02 | 2.52E-06 |
| **SAUSA300_RS05565** | srtB | class B sortase | Transport (Iron) | -1.17 | 9.13E-34 |
| **SAUSA300_RS13980** | x | GTP-binding protein | Transport (Iron) | -3.23 | 1.60E-20 |
| **SAUSA300_RS13985** | x | ferrous iron transporter B | Transport (Iron) | -3.60 | 1.21E-13 |
| **SAUSA300_RS13990** | x | NAD(P)-binding domain-containing protein | Transport (Iron) | -3.60 | 1.21E-13 |
| **SAUSA300_RS12980** | adcA | zinc ABC transporter substrate-binding lipoprotein | Transport (Metal) | -1.55 | 4.70E-12 |
| **SAUSA300_RS13350** | cntA | staphylopine-dependent metal ABC transporter substrate-binding protein | Transport (Metal) | -1.04 | 4.18E-28 |
| **SAUSA300_RS13345** | cntB | ABC transporter permease | Transport (Metal) | -1.10 | 7.58E-87 |
| **SAUSA300_RS13340** | cntC | ABC transporter permease | Transport (Metal) | -1.10 | 7.58E-87 |
| **SAUSA300_RS13335** | cntD | ABC transporter ATP-binding protein | Transport (Metal) | -1.10 | 7.58E-87 |
| **SAUSA300_RS13330** | cntF | ABC transporter ATP-binding protein | Transport (Metal) | -1.10 | 7.58E-87 |
| **SAUSA300_RS13360** | cntL | D-histidine (S)-2-aminobutanoyltransferase | Transport (Metal) | -1.51 | 1.75E-19 |
| **SAUSA300_RS13355** | cntM | staphylopine dehydrogenase | Transport (Metal) | -1.51 | 1.75E-19 |
| **SAUSA300_RS11200** | kdpD | sensor histidine kinase | Transport (Metal) | -1.05 | 4.75E-04 |

| | | | | | |
|---|---|---|---|---|---|
| **SAUSA300_RS11205** | kdpE | response regulator transcription factor | Transport (Metal) | -1.05 | 4.75E-04 |
| **SAUSA300_RS11195** | kdpF | K(+)-transporting ATPase subunit F | Transport (Metal) | -2.63 | 6.64E-04 |
| **SAUSA300_RS02270** | zagA | GTP-binding protein | Transport (Metal) | -1.24 | 2.02E-03 |
| **SAUSA300_RS00755** | phnC | phosphonate ABC transporter ATP-binding protein | Transport (Phosphate) | -1.21 | 5.64E-05 |
| **SAUSA300_RS13920** | x | PTS transporter subunit IIC | Transport (Phosphate) | -1.19 | 1.41E-06 |
| **SAUSA300_RS00130** | AdsA | LPXTG-anchored adenosine synthase | Virulence | -1.00 | 1.22E-16 |
| **SAUSA300_RS10530** | chp | chemotaxis-inhibiting protein CHIPS | Virulence | -1.22 | 1.83E-05 |
| **SAUSA300_RS05680** | flr | formyl peptide receptor-like 1 inhibitory protein | Virulence | -1.41 | 1.01E-19 |
| **SAUSA300_RS10930** | hld | delta-lysin family phenol-soluble modulin | Virulence | -2.06 | 5.69E-09 |
| **SAUSA300_RS04185** | nuc | thermonuclease family protein | Virulence | -1.41 | 4.78E-21 |
| **SAUSA300_RS00515** | plc | phosphatidylinositol-specific phospholipase C | Virulence | -1.35 | 4.28E-06 |
| **SAUSA300_RS15740** | *psmα1* | phenol-soluble modulin PSM-alpha-1 | Virulence | -3.97 | 9.49E-09 |
| **SAUSA300_RS15735** | *psmα2* | phenol-soluble modulin PSM-alpha-2 | Virulence | -4.01 | 8.32E-09 |
| **SAUSA300_RS15090** | *psmα3* | phenol-soluble modulin PSM-alpha-3 | Virulence | -4.06 | 5.68E-09 |
| **SAUSA300_RS15730** | *psmα4* | phenol-soluble modulin PSM-alpha-4 | Virulence | -4.04 | 5.17E-09 |
| **SAUSA300_RS05790** | *psmβ1* | beta-class phenol-soluble modulin | Virulence | -3.16 | 6.47E-10 |
| **SAUSA300_RS05795** | *psmβ2* | beta-class phenol-soluble modulin | Virulence | -3.18 | 1.28E-09 |
| **SAUSA300_RS10525** | scn | complement inhibitor SCIN-A | Virulence | -1.49 | 5.86E-28 |
| **SAUSA300_RS10340** | scpA | cysteine protease staphopain A | Virulence | -1.34 | 3.11E-08 |
| **SAUSA300_RS09615** | splB | serine protease | Virulence | -1.26 | 2.32E-10 |
| **SAUSA300_RS10345** | x | staphostatin A | Virulence | -1.02 | 1.38E-05 |
